# An intracellular phosphorus-starvation signal activates the PhoB/PhoR two-component system in *Salmonella enterica*

**DOI:** 10.1101/2023.03.23.533958

**Authors:** Roberto E. Bruna, Christopher G. Kendra, Mauricio H. Pontes

## Abstract

Bacteria acquire P primarily as inorganic orthophosphate (Pi, PO_4_^3-^). Once internalized, Pi is rapidly assimilated into biomass during the synthesis of ATP. Because Pi is essential, but excessive ATP is toxic, the acquisition of environmental Pi is tightly regulated. In the bacterium *Salmonella enterica* (*Salmonella*), growth in Pi-limiting environments activates the membrane sensor histidine kinase PhoR, leading to the phosphorylation of its cognate transcriptional regulator PhoB and subsequent transcription of genes involved in adaptations to low Pi. Pi limitation is thought to promote PhoR kinase activity by altering the conformation of a membrane signaling complex comprised by PhoR, the multicomponent Pi transporter system PstSACB and the regulatory protein PhoU. However, the identity of the low Pi signal and how it controls PhoR activity remain unknown. Here we characterize the PhoB-dependent and independent transcriptional changes elicited by *Salmonella* in response to P starvation, and identify PhoB-independent genes that are required for the utilization of several organic-P sources. We use this knowledge to identify the cellular compartment where the PhoR signaling complex senses the Pi-limiting signal. We demonstrate that the PhoB and PhoR signal transduction proteins can be maintained in an inactive state even when *Salmonella* is grown in media lacking Pi. Our results establish that PhoR activity is controlled by an intracellular signal resulting from P insufficiency.

## Introduction

Phosphorous (P) is essential for life. As an integral component of lipids, nucleic acids and nucleotides, this element accounts for as much as 5.5% of the dry weight of actively growing cells (1, 2). In nature, the vast majority of P exists as inorganic orthophosphate (Pi, PO_4_^3-^), either as hydrated phosphate minerals or soluble phosphate ions. Pi is the preferred P source of most living cells, and is normally assimilated into biomass during the synthesis of ATP. ATP functions as the main Pi donor molecule and the primary source of high energy phosphoanhydride bonds. Cells regulate Pi uptake to obtain adequate supplies of this nutrient and avoid toxicity resulting from accumulation of intracellular Pi and uncontrollable ATP synthesis (3–7).

In the bacterial species *Escherichia coli* and *Salmonella enterica* (*Salmonella*), Pi is acquired primarily via the canonical Pi transporters PitA and PstSCAB. PitA is a housekeeping, low affinity transporter, which is responsible for Pi uptake under most growth conditions (3, 4, 8– 11). PstSCAB is an inducible, high-affinity, ATP-dependent Pi transporter comprised of four proteins: An extra-cytoplasmic Pi-binding protein (PstS), two transmembrane proteins that form a Pi channel (PstA and PstC), and a cytoplasmic ATPase component (PstB) that couples ATP hydrolysis with Pi uptake (4, 6). These proteins are encoded in the *pstSCAB-phoU* operon, which is transcriptionally activated by the PhoB/PhoR two-component signal transduction system (4, 6, 12).

PhoR is a membrane-bound bifunctional histidine kinase/phosphatase and PhoB is its cognate transcriptional response regulator. This two-component signal transduction system has been primarily studied in the context of Pi starvation. Growth in low environmental Pi promotes PhoR kinase activity. PhoR auto-phosphorylates and subsequently transfers the phosphoryl group to PhoB (13, 14). Phosphorylated PhoB binds to DNA sequences, known as Pho boxes, at the promoter regions of target genes and stimulates their transcription (15–19). Members of the PhoB-regulon with known functions have been historically characterized in *E. coli* and include those directly involved in Pi signaling and transport (*pstSCAB*, *phoBR, phoE*) and in the extraction of Pi from alternative organic P-sources (e.g., *phn* and *ugp* genes) (5, 6).

In prototypical two-component signal transduction systems, sensor histidine kinases often harbor an extra-cytoplasmic domain that participates in the recognition of a cognate regulatory molecule or signal. However, PhoR lacks a sizable extra-cytoplasmic domain, suggesting that this histidine kinase does not directly respond to an extra-cytoplasmic ligand (14, 20, 21). In fact, the ability of PhoR to respond to Pi levels requires both the PstSCAB transporter and the peripheral membrane protein PhoU. PhoU physically interacts with PhoR and the PstB component of the PstSCAB system (22–24). Growth under P-limiting conditions is thought to alter these physical interactions, fostering the kinase state of PhoR, phosphorylation of PhoB and the subsequent activation of the PhoB regulon (24–26). Whereas the activity of PhoR is influenced by interactions with PhoU and components of the PstSCAB transporter, the identity and location of the regulatory signal, and the mechanism by which it controls PhoR activity remain poorly understood. Nevertheless, it is commonly accepted that low extra-cytoplasmic Pi promotes the kinase state of PhoR (5, 22, 24–30).

We previously established that the PhoB/PhoR systems of wild-type *E. coli* and *Salmonella* are activated when these organisms are grown in media containing abundant Pi while experiencing translational arrest resulting from either cytoplasmic Mg^2+^ starvation, or treatment with antibiotics that inhibit protein synthesis (3, 31, 32). These results implied that this two-component system responds to an intracellular signal that is normally generated during growth in low Pi but that can be brought about by these other physiological disturbances. Here, we characterize the transcriptional response of *Salmonella* to P starvation and identify PhoB-dependent and independent genes that are required for the utilization of various organic P-sources. We use this knowledge to genetically test whether the PhoB/PhoR responds to extra-cytoplasmic Pi or cytoplasmic signal(s). We demonstrate that if *Salmonella* is provided with an alternative P source that is metabolized in a PhoB-independent manner, PhoB/PhoR remains inactive during growth in Pi-free media. Furthermore, we show that PhoB/PhoR can be activated in high Pi media in response to genetic manipulations that are expected to reduce intracellular Pi. Together, our results establish that PhoB/PhoR is activated by an intracellular signal resulting from P insufficiency (Fig. 1).

**Figure 1.**
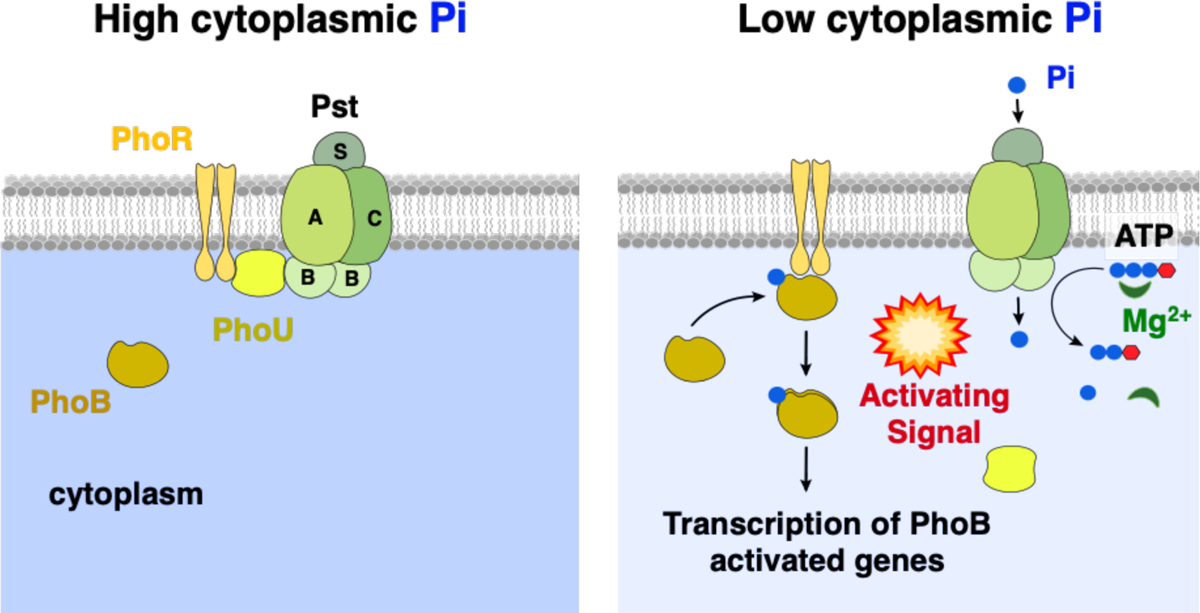
Environmental control of PhoB/PhoR. Schematics depicting the multicomponent signal complex controlling Pi-responsive PhoB/PhoR two-component system activity. Left-hand-side panel: In Pi abundant conditions, the PhoU regulatory protein interacts with and represses the activity of the Pst transport system and PhoR kinase activity. Right-hand-side panel: A decrease in cytoplasmic Pi levels alters or disrupts the PhoU interaction with Pst and PhoR. Pst begins to import extracellular Pi while PhoR phosphorylates its cognate response regulator, PhoB. PhoB-P activates the transcription of its target genes, including the *pstSCAB-phoU* operon, *phoBR*, and other genes required for Pi acquisition.

## Results

### Overview of the transcriptomic response of *Salmonella* to P starvation

When exponentially growing *Salmonella* undergoes a nutritional downshift from Pi-rich medium to medium lacking a P source, PhoB/PhoR undergoes a rapid activation surge: Fluorescence derived from transcriptional fusions between the PhoB-activated genes *phoB* or *pstS* and an unstable green fluorescence protein (*gfp*AAV) peaked at approximately 35 min following the downshift (Fig. S1A). In contrast, the activities of these fluorescent fusions did not change considerably when, instead, cells were shifted to fresh Pi-rich medium (Fig. S1A). Both the wild-type and *phoB* strains of *Salmonella* grew at similar rates, reached equivalent growth yields, and retained equal viability after 35 min in medium lacking P (Fig. S1B and S1C). Therefore, failure to activate PhoB neither impaired growth nor affected bacterial viability during P starvation (33–36).

We used RNA-seq to identify PhoB-dependent and independent transcriptional changes elicited by P starvation in *Salmonella*. We compared the transcriptomic profiles of exponentially growing wild-type and *phoB* strains at 35 min following shifts from Pi-rich medium to either medium lacking a P source or fresh Pi-rich medium (Fig. 2A). In wild-type cells, P starvation caused the upregulation of 180 genes and the downregulation of 201 genes (≥ 4-fold change, adj. P-value <0.01). In comparison, 91 and 287 genes were up- and downregulated, respectively in *phoB* cells (Fig. 2B and 2C, Table S1). In both wild-type and *phoB* strains, upregulated transcripts were overrepresented within Cluster of Orthologous Genes (COG) categories encoding proteins with unknown and general function (Fig. 2C). Prominently, P starvation caused a PhoB-independent increase in the transcription of genes within *Salmonella* pathogenicity island-2 (SPI-2) (Fig. 2C, Fig. S2 and Table S1) (36–40). This horizontally acquired chromosomal region encodes several virulence proteins, including the structural components of a type III secretion system that is required for *Salmonella* replication within mammalian macrophages (41–45).

**Figure 2.**
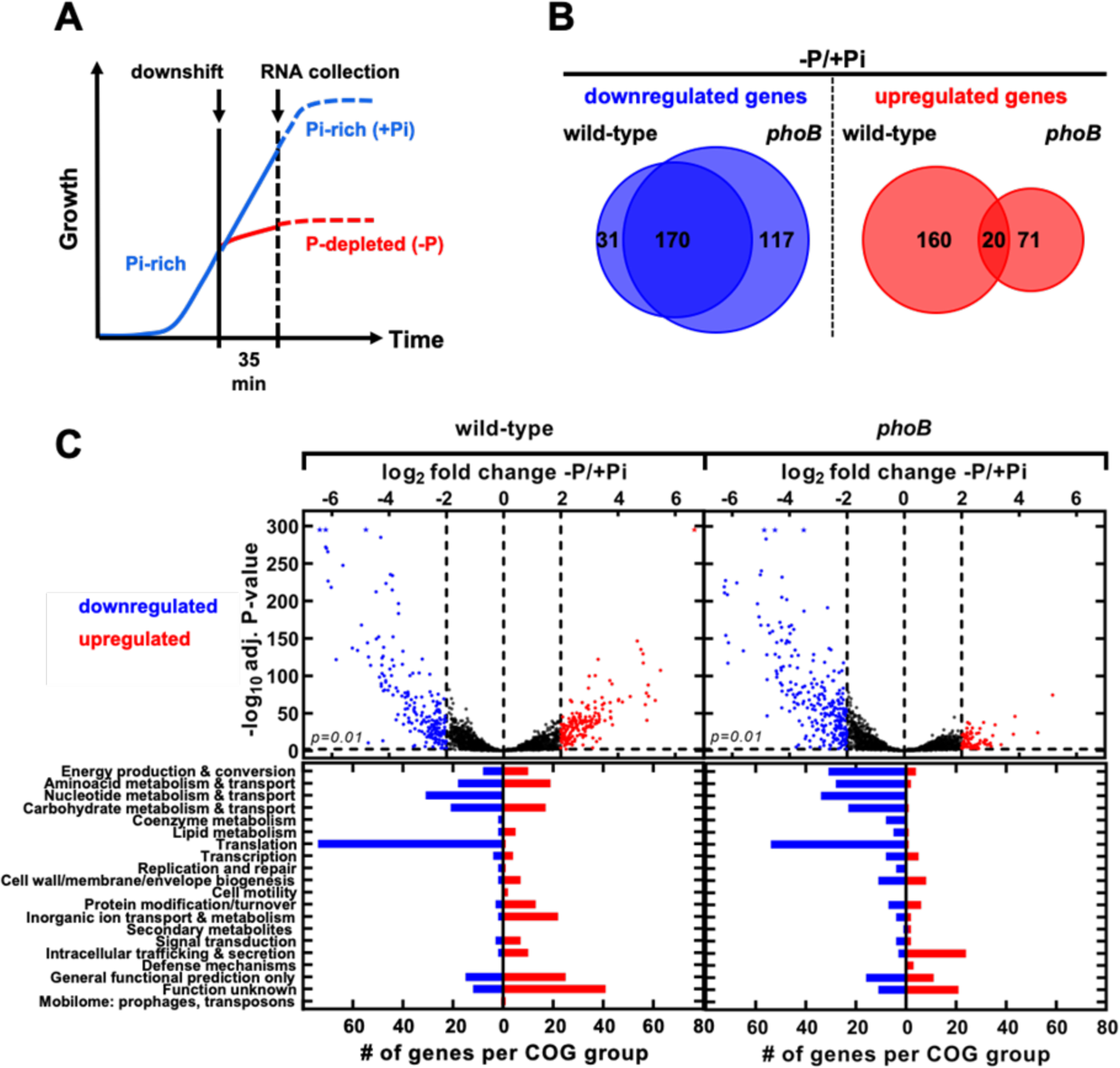
Overview of *Salmonella* transcriptomic response to P starvation. **(A)** RNA-seq experimental design. Wild-type (14028s) and *phoB* (EG9054) *Salmonella* cells were grown to mid-exponential phase in Pi-rich (+Pi, 1 mM K_2_HPO_4_) MOPS medium. At this point, cultures were washed in Pi-free MOPS medium and split in halves: one-half was inoculated in MOPS medium lacking a P source, while the second one was maintained in Pi-rich (1 mM K_2_HPO_4_) MOPS. Growth continued for additional 35 min, before proceeding to RNA extraction. Further experimental details are described in the Materials and Methods. **(B)** Quantitative Venn diagrams representing genes whose corresponding mRNAs are at least 4-fold different (log_2_fold change >2) between -P and +Pi treatments. **(C)** *(Top)* Volcano plots depicting the expression fold change versus the -log_10_ of the corresponding adjusted P-value for all genes between – P and +Pi conditions in wild-type and *phoB* mutant strains. Blue dots represent genes exhibiting a ≥ 4-fold decrease in transcripts, whereas red dots represent genes exhibiting a ≥ 4-fold increase in transcripts between conditions. Asterisks (*) represent transcripts with clipped values: logs of very small adj. p-values are represented as 295. Data correspond to DESeq2 expression analysis of 4 biological replicates using Geneious Prime software. *(Bottom)* Frequency distribution of assigned COG categories for genes exhibiting a ≥ 4-fold decrease (blue bars) or increase (red bars) in expression between -P and +Pi treatments.

P starvation downregulated 170 transcripts in both wild-type and *phoB* cells (Fig. 2B, 2C and Table S1). Downregulated RNA species were overrepresented in COG categories of proteins involved in ribosomal structure, biogenesis and translation (i.e., ribosomal proteins, ribosomal maturation factors, elongation and release factors), nucleotide transport and metabolism (i.e., purine, pyrimidine, nucleoside and nucleotide biosynthetic pathways), and amino acid transport and metabolism (i.e., polyamine biosynthesis) (Fig. 2B, 2C, S2 and Table S1). Curiously, many of these genes are transcriptionally repressed by the alarmone (p)ppGpp (46–50). During P starvation, (p)ppGpp accumulates as a result of the inhibition of the hydrolytic activity of the bifunctional (p)ppGpp synthase/hydrolase SpoT (51–53). Because (p)ppGpp promotes the transcription of SPI-2 genes (49, 54–56), the effect of P starvation on SPI-2 transcription (Fig. S2) may result from changes in SpoT’s activity.

P starvation also increased transcripts from 160 genes in the wild-type but not in the *phoB* mutant strain (Fig. 2B and 2C, Table S2; > 4-fold change, adj. p-value < 0.01). Most of these genes belong to COG categories encoding proteins involved in energy production and conversion, carbohydrate, amino acid, and inorganic ion transport and metabolism (Fig. 2C and 3). These include several members of the PhoB regulon that were previously characterized in *E. coli* (i.e., *phoBR*, *phoE*, *pstSCAB-phoU*, *ugpBAECQ*, *phnXWRSTUV*, *psiE*) (5, 6) and *Salmonella* (i.e., *apeE*, *nagA* and *nagB*) (36, 57) (Fig. 3). Because a substantial fraction of transcription start sites (TSS) have been inferred in *Salmonella* (38, 49), we used the web-based MEME Suite (58) to search for Pho boxes within or at the vicinity of the promoter regions of the aforementioned PhoB-dependent transcripts (15, 59). Putative Pho boxes were identified upstream of 28 transcriptional units (Fig. 3 and S3, Tables S3 and S4), suggesting that downstream gene(s) may be under direct PhoB control. Taken together, these results indicate that P starvation triggers a rapid transcriptional reprograming in *Salmonella*. Whereas most of the observed changes reflect PhoB-independent repression of transcripts that are often associated with cell growth (e.g., stringently repressed genes), most upregulated transcripts are PhoB-dependent.

**Figure 3.**
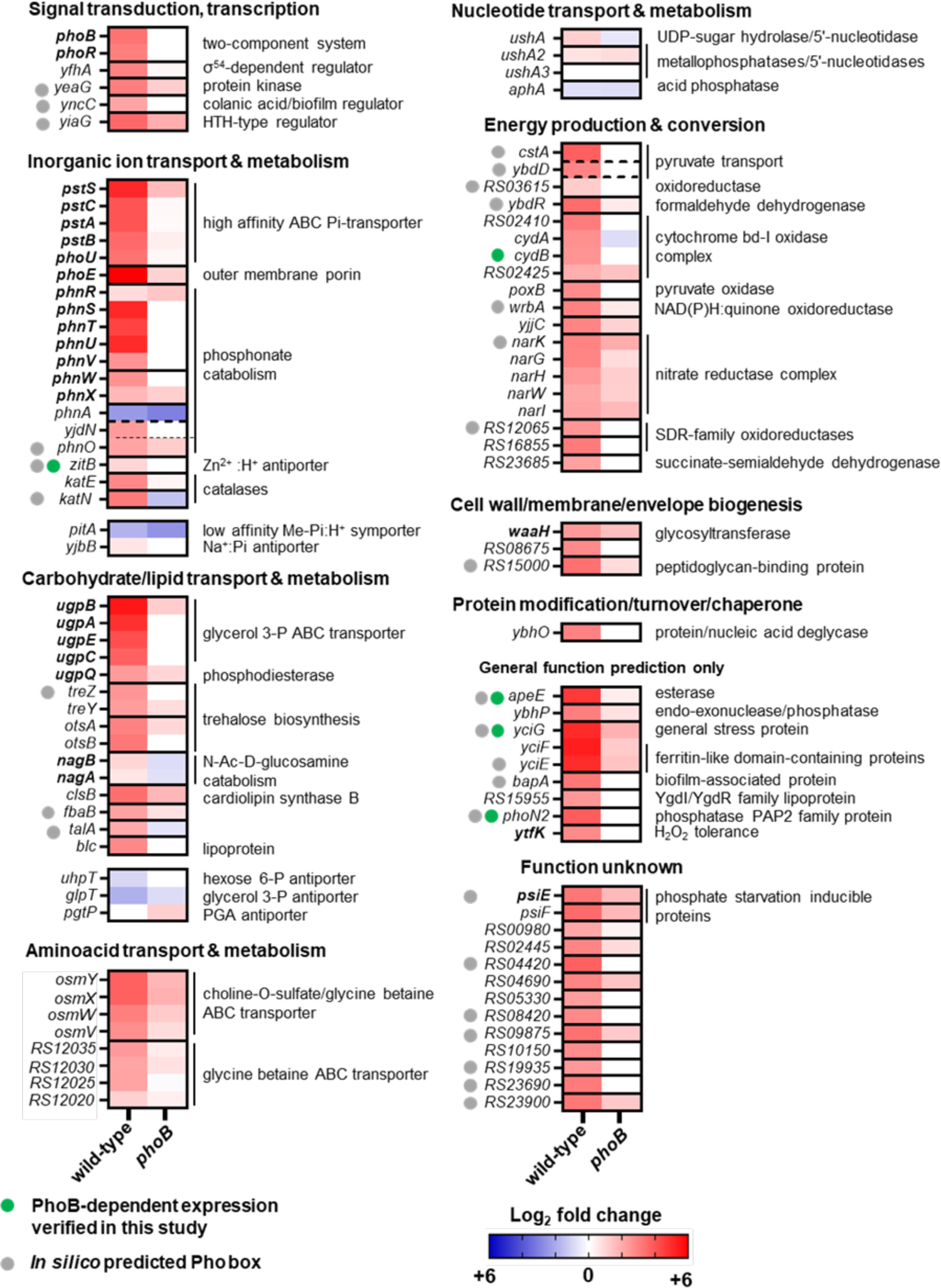
PhoB-dependent transcriptional changes during *Salmonella* P-starvation. Heatmaps depicting fold changes in transcript levels between -P and +Pi treatments for wild-type (14028s) and *phoB* (EG9054) *Salmonella*. Graphs show transcripts from genes with the highest fold change between wild-type and *phoB* during P starvation (log_2_ < −3; Tables S1 and S2). Genes are grouped by COG categories. A subgroup of PhoB-independent P-acquisition-related genes is depicted. In bold, members of the Pho regulon that have been identified in previous studies (in either *E. coli* or *Salmonella*) for which direct PhoB-regulation has been confirmed by electrophoretic mobility and/or DNA footprinting assays. Genes whose PhoB-dependent up-regulation during Pi-starvation was experimentally verified in this study (Fig. 4) are labeled with. Genes in which putative PhoB-binding motifs were identified *in silico* in this study are labeled with. Note that NCBI gene locus tags from the 14028s genome are abbreviated (e.g., locus tag *STM14_RS12020* is simply displayed as *RS12020*).

### PhoB-dependent expression of ZitB, CydB, YciG, ApeE and PhoN2 during P starvation

We sought to measure the effect of P starvation on protein levels of five PhoB-activated genes: *zitB*, *cydB*, *yciG, apeE* and *phoN2* (Fig. 3 and 4A, Table S2). To this end, we engineered a set of *Salmonella* strains, each harboring a C-terminally HA-tagged version of these genes at their native chromosomal locations. Logarithmically growing, HA-tagged wild-type and *phoB* cells were subjected to a downshift from Pi-rich medium to medium lacking P, and protein levels were assayed by Western blot. We determined that P starvation increased the amounts of ZitB-, CydB-, YciG-, ApeE- and PhoN2-HA in the wild-type but not in the *phoB* mutant strain (Fig. 4B). These results independently verify part of the transcriptional data presented above (Fig. 3 and Table S2).

**Figure 4.**
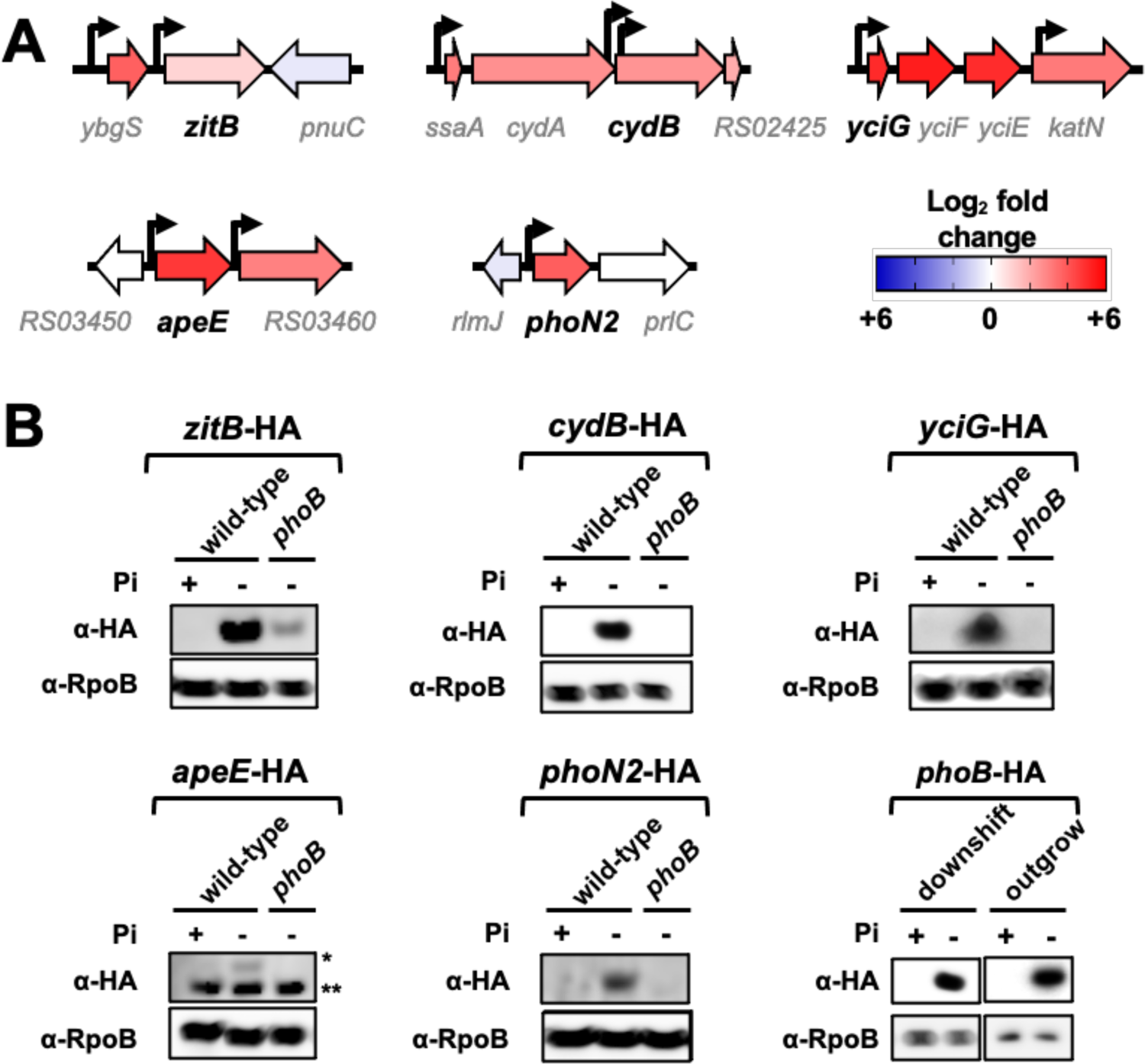
Verification of RNA-seq results. **(A)** Diagrams depicting the genomic context of *phoN2* (*STM14_RS19030*), *yciG* (*STM14_RS09540*), *apeE* (*STM14_RS03455*), *zitB* (*STM14_RS04415*) and *cydB* (*STM14_RS02420*) loci in *Salmonella*. The bent arrows represent previously mapped transcriptional start sites (38, 49). Protein-coding sequences are represented by large arrows, colored according to their fold-change expression in wild-type cells subjected to P-starvation (Fig. 3, Tables S1 and S2). Note that NCBI gene locus tags from the 14028s genome are abbreviated (e.g., locus tag *STM14_RS02425* is simply displayed as *RS02425*). **(B)** Western blot analysis of protein extracts prepared from wild-type (14028s), *phoN2*-HA (RB441), *phoB phoN2*-HA (RB447), *yciG*-HA (RB439), *phoB yciG*-HA (RB445), *apeE*-HA (RB440), *phoB apeE*-HA (RB446), *zitB*-HA (RB437), *phoB zitB*-HA (RB443), *cydB*-HA (RB438), *phoB cydB*-HA (RB444), and *phoB*-HA (MP1429). Protein extracts were obtained from cultures that were either starved for P or grown in Pi-rich medium. For PhoN2-HA and YciG-HA detection, exponentially grown cells were subjected to a 35 min-long downshift to MOPS medium lacking a P source. For ApeE-HA, ZitB-HA and CydB-HA, P starvation was achieved by outgrowing cultures in MOPS medium containing 50 µM K_2_HPO_4_ during 18 h. ApeE-HA is indicated (*); the lower band (**) is a non-specific loading control. Control cells were cultured in MOPS medium containing 1 mM K_2_HPO_4_ for the duration of the experiment. Because PhoB promotes its own transcription (112), detection of PhoB-HA was used as a P starvation control. Detection of RpoB was used as a loading control. Images are representative of three independent experiments.

### PhoB-dependent genes support the growth of *Salmonella* on 2-aminoethylphosphonate and *sn*-glycerol-3-phosphate as sole P source

Our RNA-seq data indicated that PhoB promotes the transcription of *phnSTUV*, *phnWX* and *ugpBAEC* operons in response to P deprivation (Fig. 3, 5A, 5B and Table S2). The *phnSTUV* and *phnWX* operons encode components of a PhoB-dependent phosphonatase catabolic pathway, which mediates the uptake and degradation of 2-aminoethylphosphonate (2-AP) (5, 60–62). We confirmed that wild-type bacteria grow on 2-AP as the sole P source—an ability that is impaired by deletions of either *phnWRSTUV* or *phoB* (Fig. 5A). Notably, it has been reported that *Salmonella* can also utilize phosphonoacetic acid (PAA) as sole P source (63). However, contrary to this finding, we determined that both wild-type or *phoB Salmonella* strains lack this ability (Fig. S4A-C). In fact, PAA degradation requires the relaxed specificity of phophonatases encoded in the C-P lyase catabolic pathway. While this pathway is absent in *Salmonella*, it is found in several closely related species such as *Klebsiella aerogenes* (5, 62, 64), which can grow on PAA as the sole P source (Fig. S4C).

**Figure 5.**
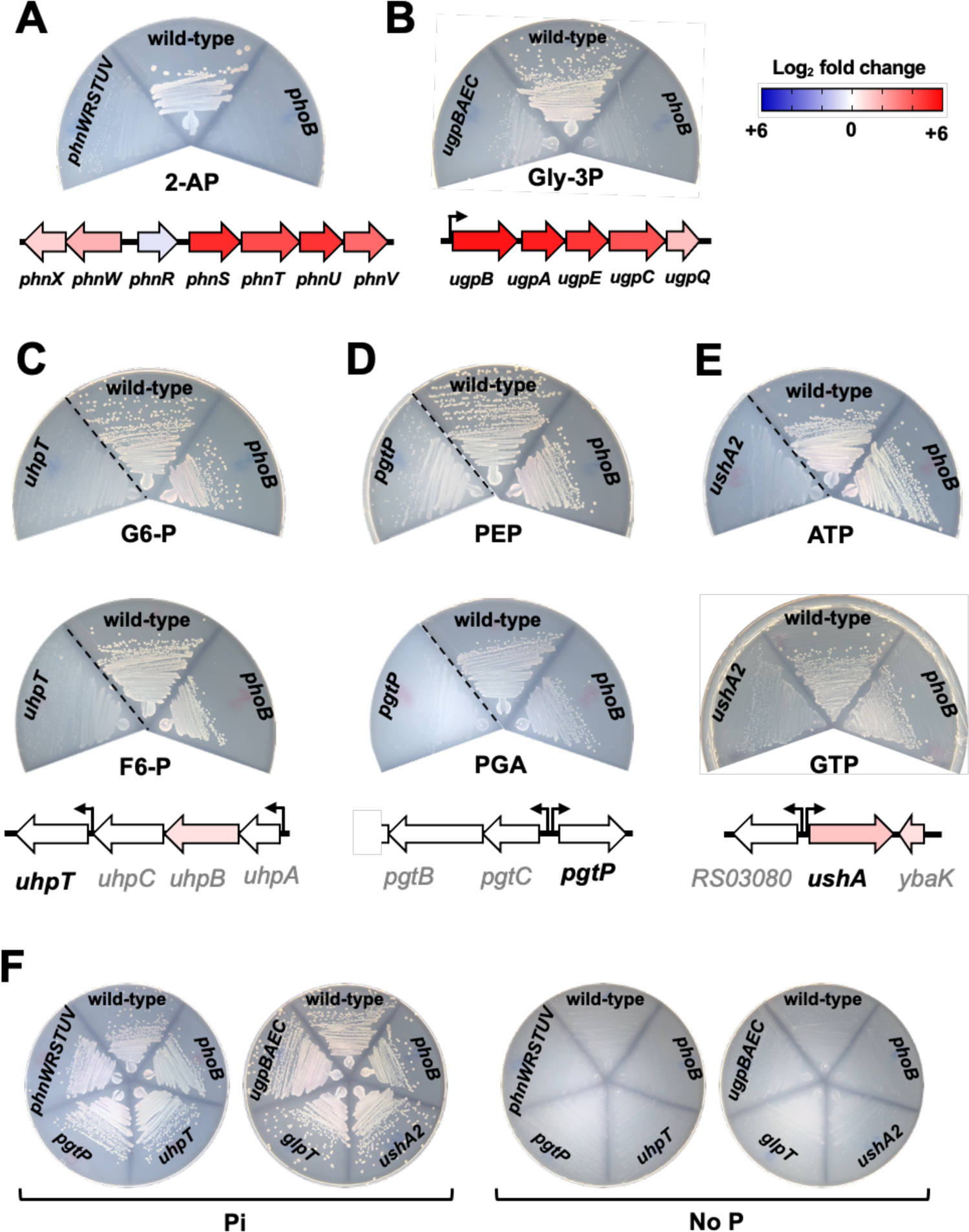
Genetic requirements of *Salmonella* for grow on different organic compounds as the sole P source. Wild-type (14028s), *phoB* (EG9054), *ugpBAEC* (MP1736), *glpT* (MP1737), *phnWRSTUV* (MP1784), *uhpT* (MP1738), *pgtP* (MP1739) and *ushA2* (MP1779) *Salmonella.* Indicated strains were grown on MOPS-glucose-noble agar media containing 1 mM of (**A**) 2-aminoethylphosphonate (2-AP), (**B**) *sn*-glycerol-3-phosphate (Gly-3P), (**C**) glucose 6-P (G6-P, top) or fructose 6-P (F6-P, bottom), (**D**) phosphoenolpyruvate (PEP, top) or 3-phosphoglyceric acid (PGA, bottom), (**E**) ATP (top) or GTP (bottom). Diagrams under the plates outline the genomic context of and expression profile of gene(s) being assayed. The bent arrows represent previously mapped transcriptional start sites (38, 49). Protein-coding sequences are depicted as large arrows that are colored according to their fold-change expression in wild-type cells subjected to P-starvation (Fig. 3, Tables S1 and S2). (**F**) Positive and negative control plates for strains tested in (A-E). Plates contain either 1 mM Pi or no P source. Plates were incubated at 37°C during 16-18 h before being imaged. Images are representative of three independent experiments. Dashed lines separate non-contiguous sections of the same plate. Note instances of residual growth due to contaminating P source(s) in the agar and, in some cases, organic P source.

In *E. coli*, the *ugpBAECQ* locus encodes a PhoB-activated *sn*-glycerol-3-P (Gly-3P) ABC transporter (UgpBAEC) and a cytoplasmic phosphodiesterase capable of releasing Pi from this substrate (UgpQ). Together, these proteins enable *E. coli* to use Gly-3P as the sole P source (65–67). Similarly, we determined that *Salmonella* can grow on Gly-3P as the sole P source, and that growth on this substrate is impaired by deletions of either *ugpBAEC* or *phoB* (Fig. 5B). Collectively, these results indicate that during P starvation, PhoB promotes the transcription of *phnSTUV*, *phnWX* and *ugpBAEC* operons, thereby allowing *Salmonella* to scavenge P from 2-AP and Gly-3P, respectively.

### PhoB-independent genes support the growth of *Salmonella* on a variety of organic-Pi sources

We wondered if *Salmonella* could utilize additional P sources in a PhoB-independent manner. This bacterium harbors homologs of three major facilitator antiporters that can uptake a cognate negatively charged organic-P substrate in counterflow with a Pi anion (68–70). GlpT transports Gly-3P (27, 71, 72), UhpT translocates hexoses-6-phosphates (73), and PgtP uptakes 2- or 3-phosphoglyceric acid (PGA) and phosphoenolpyruvate (PEP) (74, 75). It has been suggested that these antiporters can promote a net Pi uptake by asymmetric self-exchange, where two external organic-Pi molecules are exchanged against one internal organic-Pi molecule, and that other cytoplasmic anions such as sulfate can participate in the exchange reaction (5, 69). We therefore tested the abilities of wild-type and *glpT*, *uhpT* and *pgtP* mutant strains to grow on cognate organic substrates as the sole P source. We determined that GlpT is dispensable for growth on Gly-3P (Fig. S5A), which depends primarily on proteins encoded in the *ugpBAECQ* operon (Fig. 5B). In contrast, UhpT is required for the growth of *Salmonella* on medium containing either glucose-6-phosphate (G6-P) or fructose-6-phosphate (F6-P) as the sole P source. That is, an *uhpT* deletion hindered growth on G6-P or F6-P (Fig. 5C). Interestingly, *E. coli* harbors an UhpT homolog that shares 100% amino acid sequence identity with the UhpT protein from *Salmonella*. However, *E. coli* is unable to use G6-P as the sole P source (76). Similarly, a *pgtP* deletion impaired growth on either PGA or PEP as sole P source (Fig. 5D), indicating that PgtP allows *Salmonella* to utilize these substrates as the sole P source (75). Importantly, a deletion of *phoB* did not affect transcription of *uhpT* or *pgtP* (Fig. 3, Table S1), nor did it hinder the ability of *Salmonella* to grow on G6-P, F6-P, PGA or PEP as the sole P source (Fig. 5C, 5D and 5F).

P acquisition from organic sources can also be attained by the activity of periplasmic enzymes that scavenge Pi from organic substrates to enable their uptake by Pi transporters such as PitA. *Salmonella* encodes three UDP-sugar/5’-nucleotidases paralogs (*ushA*, *ushA2* and *ushA3*) as well as a periplasmic acid phosphatase (*aphA*). Whereas UDP-sugar/5’-nucleotidases can release terminal Pi from nucleoside mono-, di- or triphosphates (77, 78), AphA has a relaxed substrate specificity being able to act on a wide range of organic substrates including some nucleotides (79, 80). Transcription of these genes was not affected by PhoB during P starvation, suggesting that their expression is PhoB-independent (Fig. 3 and Table S1). Concordantly, we determined that wild-type and *phoB* strains of *Salmonella* grew robustly on plates containing either AMP, ADP, ATP or GTP as the sole P source (Fig. 5E, S5B and S5C). Whereas growth on these substrates was impaired by a deletion of *ushA2*, inactivation of either *ushA*, *ushA3* or *aphA*, did not hinder growth (Fig. 5E, S5B and S5C). Taken together, these results establish that the PhoB-independent *uhpT*, *pgtP* and *ushA2* gene products allow *Salmonella* to efficiently use G6-P/F6-P, PGA/PEP, and AMP/ADP/ATP/GTP as sole P-sources, respectively.

### Utilization of alternative Pi sources represses PhoR activity in the absence of Pi

Given that *Salmonella* can utilize a variety of organic-P substrates in a PhoB-independent manner (Fig. 3, 5, S4, S5, Table S1), we sought to formally determine whether PhoR activity is regulated by an extracellular or an intracellular signal(s). Specifically, we reasoned that if PhoR senses extra-cytoplasmic Pi, then fluorescence derived from a PhoB-dependent P*pstS-gfp* should increase in wild-type, *uhpT*, *pgtP* or *ushA2* cells when these strains are shifted from Pi-rich medium to one that lacks Pi, regardless of the availability of alternative P sources (Fig. 6). In contrast, if PhoR senses an intracellular signal that is generated during P starvation, then P*pstS-gfp* activity should remain low in wild-type cells that are shifted from Pi-rich medium to media that lacks Pi but contain an alternative organic-P source that is catabolized in a PhoB-independent fashion (e.g., G6-P, F6-P, PGA, ATP or GTP) (Fig. 6). However, in this scenario, P*pstS-gfp* fluorescence should increase in mutants that are unable to efficiently utilize these substrates, given that they are expected to experience a decrease in cytoplasmic Pi (Fig. 5 and 6) (72).

**Figure 6.**
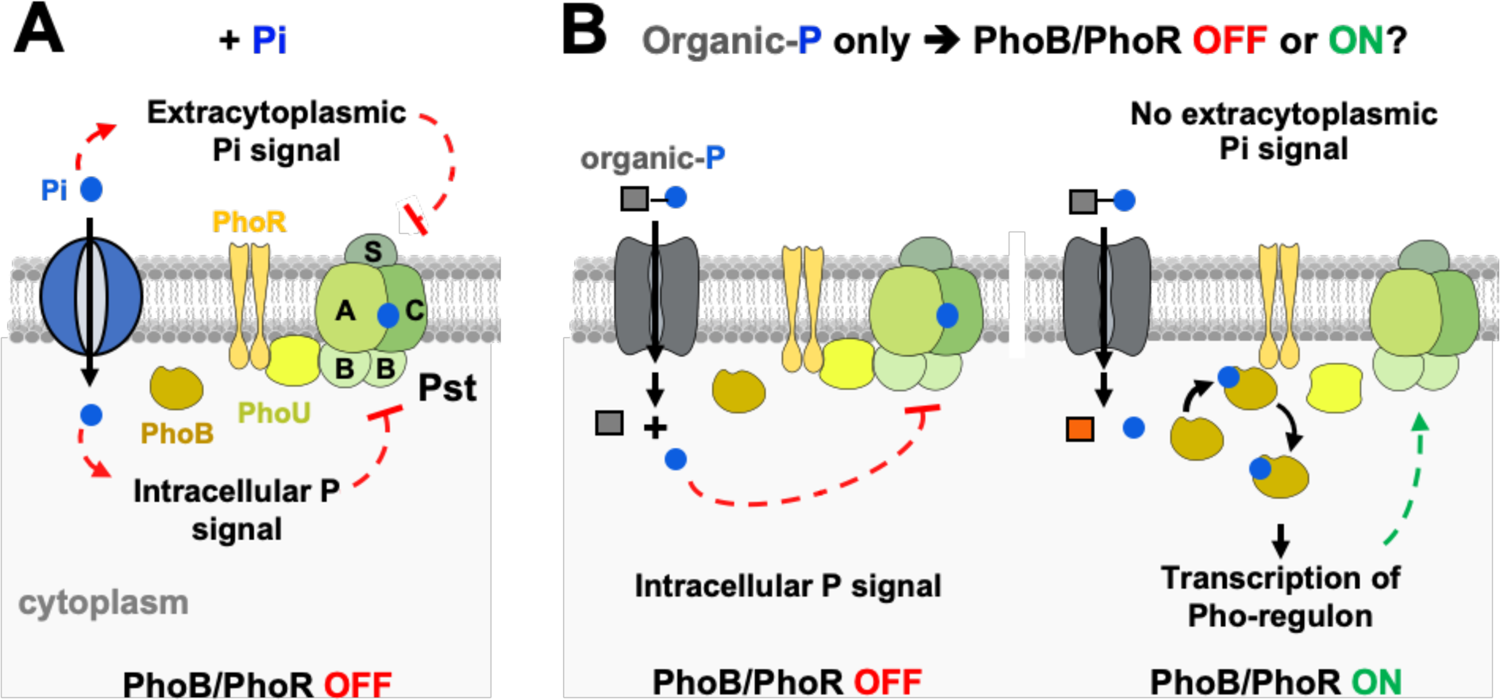
Distinguishing between extra- and intracellular sensing hypotheses. Cartoons depicting the PhoB/PhoR signaling complex (PhoR, PhoU, PstB, PstA, PstC, PstS and PhoB), a Pi transporter (blue) and an organic-P transporter (gray). (**A**) In the extracellular sensing model, Pi is sensed at the periplasmic face of the signaling complex. During growth in Pi-rich medium, high extracellular Pi signals through the Pst transporter, PhoU and/or PhoR to favor PhoR’s phosphatase conformation. In the intracellular sensing model, Pi, or a related P-sufficiency signal, is sensed at the cytoplasmic face of the signaling complex. Hence, during growth in Pi-rich medium, sufficient amounts of Pi are imported into the cytoplasm to inhibit the signaling complex and promote phosphatase state of PhoR. Classical Pi starvation experiments cannot distinguish between these two hypotheses because intra- and extracellular concentrations of Pi cannot be selectively uncoupled. (**B**) However, bacteria can be grown in medium lacking Pi and containing, instead, an organic-P substrate that is metabolized in a PhoB-independent manner. If PhoR is controlled by an intracellular signal, the concentration of intracellular P will be maintained through the PhoB-independent pathway, and the PhoB/PhoR signaling complex will remain in an OFF state. If PhoR is controlled by an extracellular signal, PhoB/PhoR will be activated.

Accordingly, we subjected exponentially growing wild-type, *uhpT*, *pgtP*, and *ushA2 Salmonella* strains to a shift from Pi-rich medium to media containing either G6-P, F6-P, PGA, ATP or GTP as the sole P source. Surprisingly, despite the lack of Pi in the growth media, P*pstS-gfp* activity remained low in wild-type cells (Fig. 7A-H). In contrast, P*pstS-gfp* fluorescence increased when the *uhpT*, *pgtP* and *ushA2* mutant strains were shifted to media containing G6-P/F6-P, PGA or ATP/GTP, respectively (Fig. 7A-H). Regardless of the genetic background, P*pstS-gfp* activity remained low following shifts to fresh Pi-rich medium and increased following shifts to P-deficient medium (Fig. 7B, 7D, 7E, 7G, 7H).

**Figure 7.**
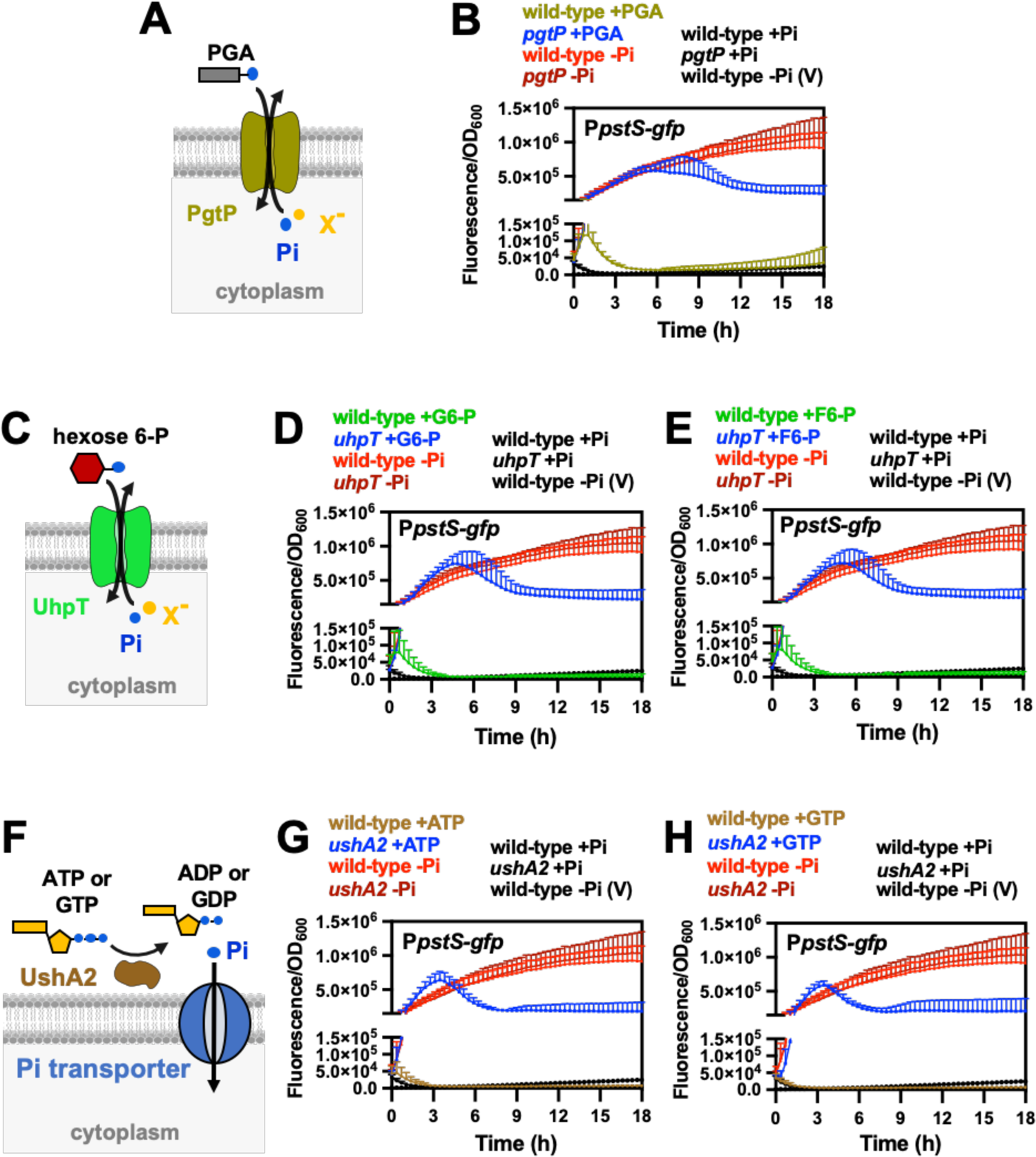
Effect of organic-P utilization on PhoB/PhoR activity. (**A**) Schematic representation of *Salmonella* PgtP protein. PgtP exchanges extracellular phosphoenolpyruvate (PEP) or 2- or 3-phosphoglycerate (PGA) with intracellular Pi or similar anions (X^-^), and promotes *Salmonella* growth on these substrates as the sole P-sources (Fig. 5 and S6C) (69, 74, 75). (**B**) Fluorescence from wild-type (14028s) and *pgtP* (MP1739) carrying the pP*pstS-gfp* or the promoterless pGFP control vector (V). Prior to fluorescent readings, cultures were grown to mid-logarithmic phase in MOPS medium containing 1 mM K_2_HPO_4_, washed in MOPS medium lacking a P source and resuspended in MOPS medium containing 1 mM PGA, 1 mM Pi or lacking a P source. (**C**) Schematic representation of *Salmonella* UhpT protein. UhpT exchanges extracellular hexose-P compounds, such as glucose 6-P (G6-P) and fructose 6-P (F6-P) with intracellular Pi or similar anions (X^-^) (69), thereby supporting the growth of *Salmonella* on G6-P or F6-P as sole P sources (Fig. 5, S6A and S6B). (**D** and **E**) Fluorescence from wild-type (14028s) and *uhpT* (MP1738) carrying pP*pstS-gfp* or pGFP (V). Fluorescent readings were conducted as described in (B), except that cultures were resuspended in medium containing either 1 mM of G6-P (D) or F6-P (E) as the sole P source. (**F**) Schematics of *Salmonella* UshA2 protein. UshA2 is a periplasmic 5’ nucleotidase capable of extracting Pi from several compounds such as ATP, ADP, AMP and GTP (Fig. 5, S5B, S5C, S6D and S6E). (**G** and **H**) Fluorescence from wild-type (14028s) and *ushA2* (MP1779) *Salmonella* carrying pP*pstS-gfp* or pGFP (V). Fluorescent readings were conducted as described in (B), except that cultures were resuspended in medium containing either 1 mM of ATP (G) or GTP (H) as the sole P source. In all cases, means ± SDs of at least three independent experiments are shown.

The increase in P*pstS-gfp* activity observed in the *uhpT*, *pgtP* and *ushA2* mutants is not a general consequence of reduced growth rate of mutant strains (Fig. S6A-E). This is because P*pstS-gfp* activity did not change when either an arginine auxotroph or a mannose catabolic mutant strain of *Salmonella* was subjected to growth arrest when shifted from medium containing arginine and glucose to either medium lacking arginine or containing mannose as the sole carbon source (Fig. S6F-G). Taken together, these results suggest that the activation state of PhoR is controlled by an intracellular signal resulting from P starvation.

### Transcriptional rewiring of PhoB-dependent *ugpBAECQ* allows PhoR repressesion by Gly-3P metabolism

The results depicted in Fig. 7 strongly suggested that, rather than extracytoplasmic Pi, PhoR is repressed by the availability of intracellular Pi or a ligand generated during Pi metabolism. To further test this notion, we constructed a *Salmonella* strain in which the PhoB-controlled promoter upstream of the *ugpBAECQ* operon (Fig. 3, 5B and Supplementary Materials) was replaced by a *tetA* promoter and a divergently transcribed *tetA* repressor encoded by the *tetR* gene (*tetRA-ugpBAECQ*; Fig. 8A). In this strain, *ugpBAECQ* is transcribed, and Gly-3P is metabolized, in response to anhydrotetracycline (aTc) instead of low Pi (Fig. 8A and 8B). Hence, if PhoR responds to an intracellular signal, then its kinase activity should be damped when *tetRA-ugpBAECQ* harboring strains are grown in medium containing aTc and Gly-3P as the sole P source. However, PhoR kinase activity should increase if aTc is removed.

**Figure 8.**
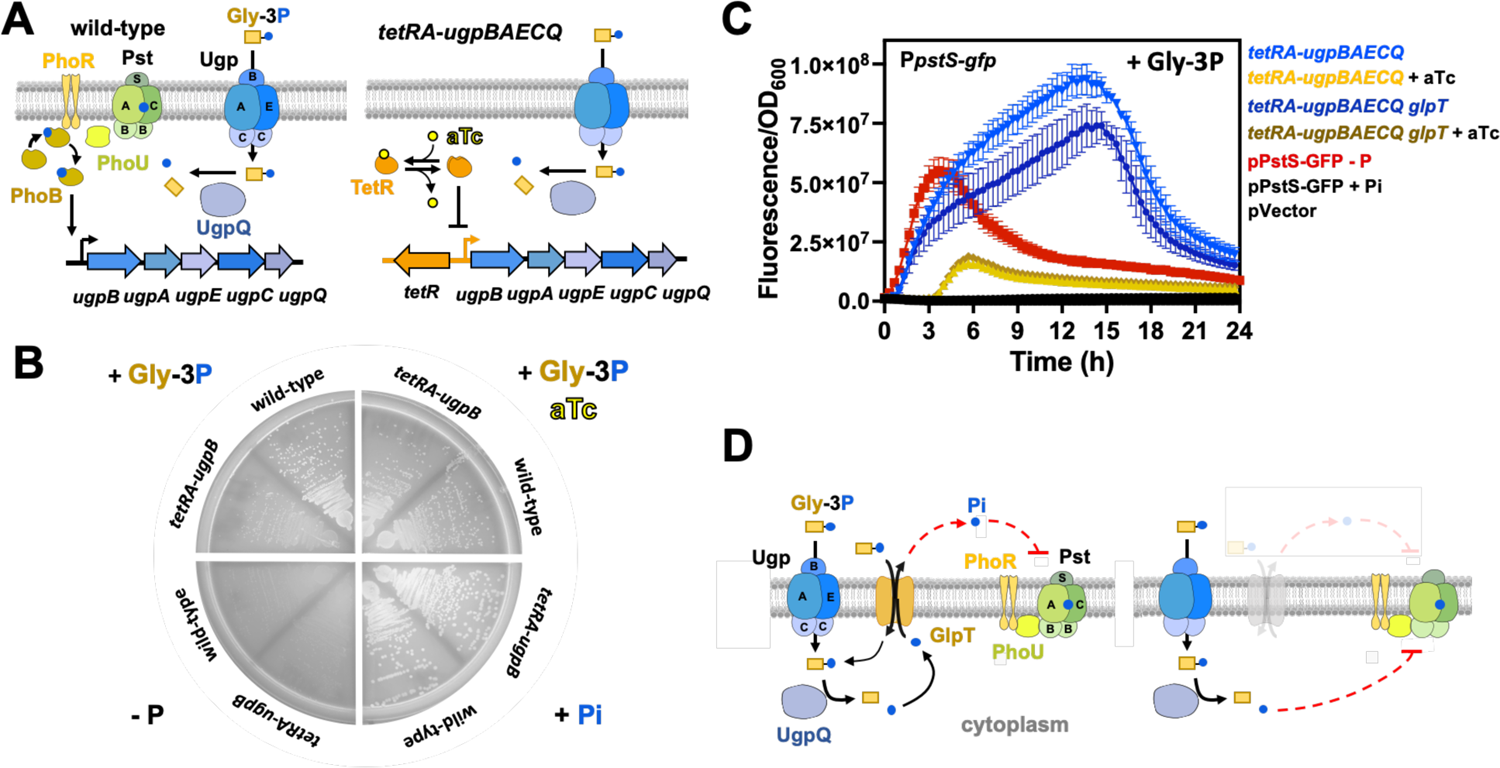
Metabolism of Gly-3P quenches PhoR activity in medium lacking Pi. **(A)** Left-hand-side panel: Cartoon representations depicting the native regulatory circuit whereby PhoB/PhoR controls the transcription of the *ugpBAECQ* operon. Right-hand-side panel: Cartoon representations depicting the engineered regulatory circuit whereby TetR controls the transcription of the *ugpBAECQ* operon in response to anhydrotetracycline (aTc). **(B)** Growth of wild-type (14028s) and *tetRA-ugpBAECQ* (MP2133) *Salmonella* strains on MOPS-glucose-noble agar plates either lacking P (-P), or containing 1 mM of Gly-3P (+Gly-3P), 1 mM of Gly-3P and 0.5µg/ml of aTc (+Gly-3P aTc), or 1 mM K_2_HPO_4_ (+Pi). Plates were incubated at 37°C for 16-18 h before being imaged. **(C)** Fluorescence from *tetRA-ugpBAECQ* (MP2133) and *tetRA-ugpBAECQ glpT* (MP2134) carrying pP*pstS-gfp* or pGFP (pVector). Strains were grown overnight in MOPS-glucose containing 2 mM K_2_HPO_4_ and the presence of absence of 0.5µg/ml of aTc. Cells were washed with MOPS-glucose lacking a P source and inoculated (1:100) into fresh MOPS-glucose containing 1 mM of Gly-3P and the presence or absence of 0.5µg/ml of aTc, and green fluorescence was monitored. Fluorescence of *tetRA-ugpBAECQ* (MP2133) harboring pP*pstS-gfp* or pVector and grown in medium lacking a P source or containing 1 mM K_2_HPO_4_ is also shown. Means ± SDs of at least three independent experiments are shown. **(D)** Left-hand-side panel: Cartoon representations depicting Gly-3P importation by the UgpBAEC transport and the release of Pi by UgpQ. GlpT can potentially exchange extracellular Gly-3P with intracellular Pi (72). Exported Pi can, presumably, repress PhoR from the periplasmic face of the signaling complex (27). Right-hand-side panel: In the absence of GlpT, Pi released by UgpQ remains in the cytoplasm to support the growth of bacteira.

Acordingly, we grew a *tetRA-ugpBAECQ* strain in medium containing Pi, in the presence or absence of aTc, and subsequently shifted the cells to the same medium containing Gly-3P as the sole P source. In cultures grown in the absence of aTc, the fluorescene derived from the PhoB-dependent P*pstS-gfp* reporter began to increase immediately after shifting cells from Pi to Gly-3P, reaching approximately 100-fold above background levels after 12 h (Fig. 8C). In contrast, for cultures grown in the presence of aTc, P*pstS-gfp*-derived fluorescence remained at background levels for the initial 4 h, subsequently increasing to approximately 10-fold above background levels after 6 h of growth (Fig. 8C). Importantly, this phenotype was recapitulated in a *tetRA-ugpBAECQ glpT* strain (Fig. 8C), that lacks the GlpT transporter and is, therefore, unable to mediate the exchange between intracellular Pi and extracellular Gly-3P (Fig. 8D). This result rules out the possibility that Pi extracted from Gly-3P is exchanged with extracytoplasmic Gly-3P to inhibit the PhoR signaling complex from the periplasmic face of the signaling complex (Fig. 8D) (27, 72). As expected P*pstS-gfp* fluorescence remained at background levels following shifts to fresh Pi-rich medium and increased following shifts to P-deficient medium (Fig. 8C). Taken together, these results establish that the kinase activity of PhoR is stimulated when cells cannot maintain sufficient levels of intracellular P because they are unable to efficiently import and metabolize Gly-3P.

### Pi transporters influence the activation state of *Salmonella* PhoR

If a cytoplasmic P starvation signal controls PhoR activity (Fig. 6-8), and P is assimilated primarily as Pi (4), then perturbations in intracellular Pi pools should also affect the activation state of this sensor kinase. We tested this prediction in a number of independent ways. First, we hypothesized that the deletion of the housekeeping Pi transporter *pitA* (Fig. 9A) would lower intracellular Pi during growth in Pi-rich medium and, as a result, increase P*pstS-gfp* reporter activity. Accordingly, deletion of *pitA* resulted in a small, but reproducible rise in P*pstS-gfp* fluorescence during growth in either complex (81) or defined Pi-rich media (Fig. 9B-C, S7). This phenotype was reversed by ectopic expression of *pitA* from a plasmid, but not by the empty plasmid control (Fig. 9B). Importantly, the increased P*pstS-gfp* activity resulted from the activation of PhoB by decreased intracellular Pi levels because P*pstS-gfp* fluorescence was abolished by either a *phoB* deletion or the ectopic expression of *pho89*, a phosphate transporter from the budding yeast *Saccharomyces cerevisiae* (82, 83) (Fig. 9A-B), without perturbing bacterial growth rate (Fig. 9D).

**Figure 9.**
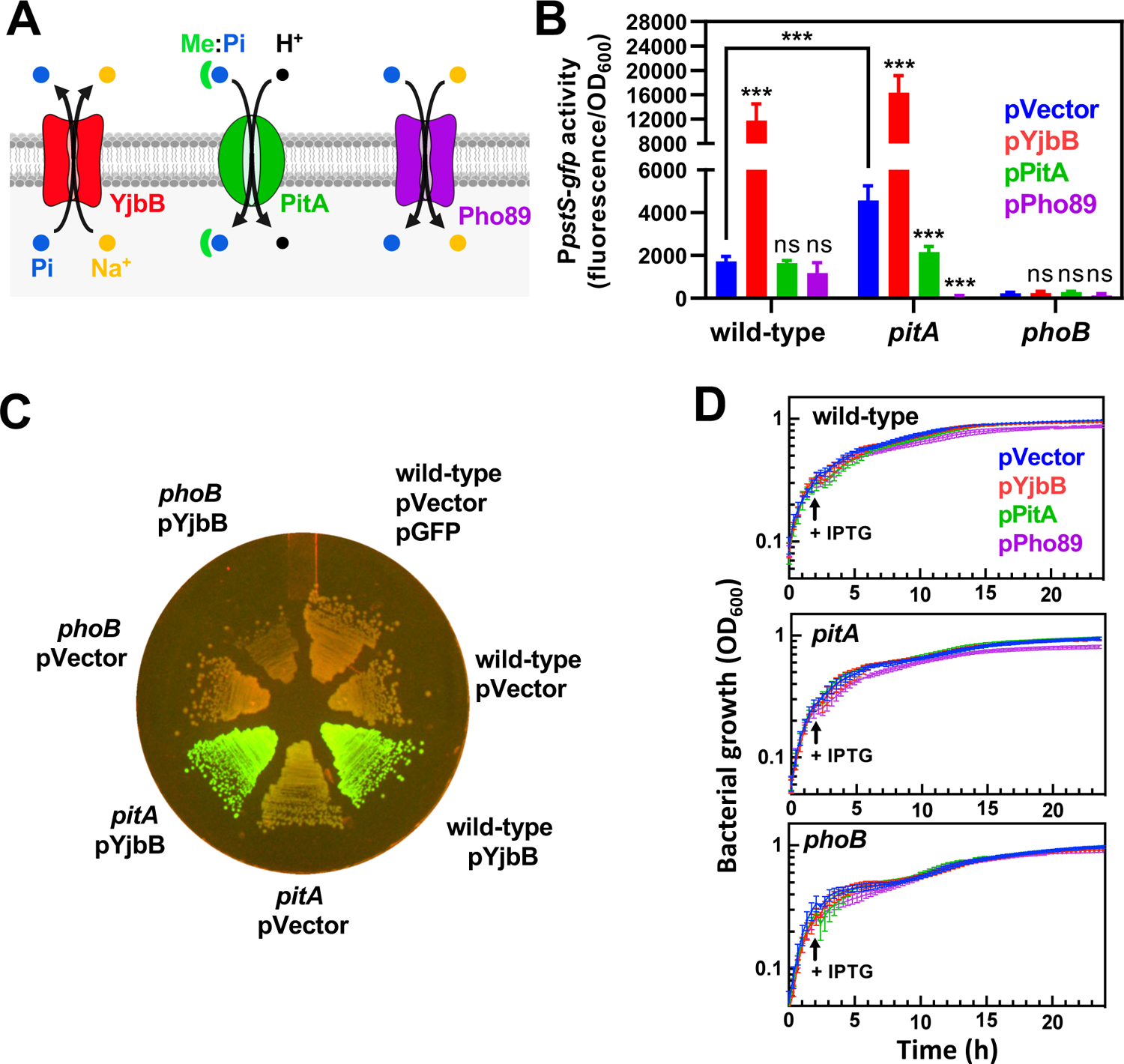
The activity of Pi importers and exporters modulates PhoB/PhoR signaling in opposite manners. **(A)** Cartoon representations of Pi transporter systems used in the described experiments. YjbB is a Na^+^/Pi co-transporter which has been shown to promote Pi export (84). PitA uses the proton motive force to transport soluble, neutral divalent metal:phosphate complexes (Me:Pi) into the cytoplasm. Pho89 is a high-affinity Na^+^/Pi co-transporter from the yeast *Saccharomyces cerevisiae*. **(B)** Fluorescence wild-type (14028s), *pitA* (MP1251), *phoB* (EG9054) *Salmonella* carrying pP*pstS*-*gfp* and either pUHE-21 (pVector), pUHE-YjbB (pYjbB), pUHE-PitA (pPitA) or pUHE-Pho89 (pPho89). Experiments were conducted in LB broth. The expression of YjbB, PitA and Pho89 were induced after 2 h of growth by the addition of 250 µM of isopropyl β-d-1-thiogalactopyranoside (IPTG). Normalized GFP values (fluorescence/OD_600_) were determined after 24 h of growth. Means ± SDs of four independent experiments are shown. ***P < 0.001; ns, no significant difference. Two-way ANOVA calculated with Dunnett multiple-comparison test. Unless indicated otherwise, statistical comparisons were made against control (pVector) samples for each of the genotypes assayed. **(C)** Fluorescence derived from wild-type (14028s), *pitA* (MP1251) and *phoB* (EG9054) *Salmonella* carrying plasmids pP*pstS-gfp* or pGFP and either pVector (pUHE-21) or pYjbB (pUHE-YjbB). Cells were grown on LB-agar plates containing 100 µM of IPTG to induce YjbB expression. Plates were incubated at 37°C during 16-18 h. Images are representative of three independent experiments. **(D)** Growth of strains depicted in (B).

Similarly, we reasoned that expression of YjbB, a Pi exporter (Fig. 9A) (84, 85), <u>would</u> stimulate PhoR kinase activity by decreasing the concentrations of cytoplasmic Pi. As hypothesized, ectopic YjbB expression resulted in 6.9- and 3.2-fold increases in P*pstS-gfp* activity in wild-type and *pitA* cells, respectively, in comparison with isogenic strains harboring the empty vector control (Fig. 9B and 9C). These increases in P*pstS-gfp* fluorescence occurred in Pi-rich medium (81) and were abolished by a *phoB* deletion (Fig. 9B and 9C). Furthermore, the effect of YjbB on *pstS-gfp* transcription occurred independent of growth rate, as *Salmonella* strains ectopically expressing YjbB grew similarly to the control strains carrying the empty vector (Fig. 9D). Altogether, the results presented in this section demonstrate that the activation state of PhoB/PhoR can be influenced by the expression of proteins that affect the concentration of cytoplasmic Pi, even when *Salmonella* is grown in high Pi environments.

## Discussion

In the current study, we define PhoB-dependent and independent transcriptional changes elicited by *Salmonella* in response to P starvation, inferring the identity of transcripts that may be under direct PhoB-control. We confirm that PhoB promotes the transcription of genes required for the importation and utilization of 2-AP and Gly-3P as the sole P sources (60, 61). Contrary to a previous report (63), we establish that *Salmonella* is unable to utilize PAA as the sole P source. We show that the PhoB-independent proteins UhpT, PgtP and UshA2 are required for the utilization of G6-P/F6-P, PGA/PEP, and ATP/ADP/AMP/GTP as the sole P source, respectively. Finally, we demonstrate that PhoR is activated by a cytoplasmic signal that is generated when bacteria are starved for P (Fig. 1).

The physiological response to P starvation has often been studied in an experimental context where bacteria are starved for P after being propagated in media containing excessive Pi. In this framework, cells experience a decrease in both extra-cytoplasmic and cytoplasmic Pi, which precipitates a molecular response aimed at restoring levels of cytoplasmic Pi (4, 6, 72, 86). In *Salmonella*, this response involves two broad patterns of transcriptional reprogramming (Fig. 2, S2 and Table S1). On the one hand, the lack of P promotes the PhoB-independent repression of several transcripts from genes that are associated with bacterial growth and replication. Most notably among these are genes encoding proteins involved in translation, including ribosomal proteins, elongation and release factors, and enzymes required for the biosynthesis of nucleotide and polyamines (87–90). Because nucleotides and rRNA are the largest intracellular reservoirs of assimilated P (3, 4, 32), inhibiting the transcription of these genes may prevent wasteful allocation of resources towards cellular components that require large quantities of P. Complementarily, the degradation of existing ribosomes induced by P starvation (91, 92), may replenish the pools of cytoplasmic nucleotides that are needed to sustain a transcriptional response to this stress.

On the other hand, P deficiency stimulates the transcription of a substantial number of PhoB-dependent genes (Fig. 2, 3, and 4, Tables S1 and S2). Whereas these genes are primarily involved in the scavenging of environmental P and the retrieval of assimilated P from cellular components (5, 6, 57, 66, 71, 93), they may also help cells to cope with secondary stresses. For instance, P starvation will lead to growth arrest (Fig. S6). However, under aerobic conditions, cells will continue to import and metabolize carbon, thereby generating endogenous reactive oxygen species that may damage cellular components (94–97). In *Salmonella*, PhoB promotes the transcription of the catalases encoded by the *katE* and *katN* genes, as well as a dedicated cytochrome bd terminal oxidase encoded by the *cydB* gene (located within the STM14_*RS02410-cydA*-*cydB*-*RS02415* operon). Whereas CydB may reduce endogenous hydrogen peroxide generation by improving oxygen consumption (96), the catalases detoxify any hydrogen peroxide that may be produced from oxygen that escapes the respiratory chain.

The PhoB-mediated response to P starvation begins with the detection of a low P signal by the PhoR signaling complex at the membrane. In this complex, PhoR physically interacts with the PstB component of the PstSCAB Pi transporter via the PhoU protein (24). Deletion of *phoU*, *pstA*, *pstB*, *pstS* or *pstSCAB-phoU* foster the kinase state of PhoR and the transcription of PhoB-dependent genes even when cells are grown in Pi-rich conditions (5, 24, 98). Whereas this indicates that protein interactions that are formed in Pi-rich conditions maintain PhoR in an inactive state, and that Pi levels may be sensed by components of the complex other than PhoR, it also suggests a way in which this complex may sense low Pi. That is, PhoR activity could be controlled by conformations adopted by the other components of the signaling complex in response to the binding of extra-cytoplasmic Pi to PstSCAB transport components (5, 22, 24–30). However, because Pi starvation regimens that are often used to study this two-component system decrease the intracellular concentration of Pi (72, 86), the above model cannot rule out the possibility that PhoR is activated by a signal sensed at the cytoplasmic face of the complex.

A number of previous studies suggest that the PhoR signaling complex may respond to an intracellular signal. Whereas inactivation of *pstS* promotes the kinase state of PhoR during growth of *E. coli* in Pi-rich medium, ectopic overexpression of phosphate transporters PitA or PitB represses PhoR kinase activity. This implies that an elevation in cytoplasmic Pi can compensate for the defect conferred by the *pstS* deletion on the signaling complex (81, 99). In the case of certain purine auxotrophic strains of *E. coli*, conditions which cause an expansion of adenine nucleotides activate PhoR during growth in otherwise Pi-rich medium (100). Similarly, treatment of *E. coli* or *Salmonella* with translation inhibitors in Pi-rich medium causes the expansion of ATP pools and promotes PhoR kinase activity (32). In these two last cases, activation likely results from a reduction in pools of free cytoplasmic Pi that are drained as a result of ATP accumulation. This is because PhoR remains active when translation-arrested cells hydrolyze excess ATP (32) or accumulate Pi alongside elevated ATP levels (3). Furthermore, under normal physiological conditions, (p)ppGpp production resulting from P starvation is expected to cause a reduction in the pools of nucleotides (Fig. 2C, S2 and Table 3) (49, 51, 52, 101).

We now provide multiple pieces of genetic evidence demonstrating that PhoB/PhoR signal transduction originates from within cells. This notion is supported by three main experimental results. First, the protein products encoded by *uhpT, pgtP* and *ushA2* allow *Salmonella* efficiently utilize Pi groups from organic substrates in a manner that is independent from both PhoB and Pi availability (Fig. 3, 5, 6, 7 and S5, S6) (75). Growth on these substrates as sole P-source supplies Pi to the cytoplasm and maintains PhoR in an inactive state. Second, the proteins encoded by the PhoB-activated *ugpBAECQ* operon allows the importation and extraction of Pi from cytoplasmic Gly-3P (Fig. 3, 5B, Table S2) (5, 72). When this operon is placed under the control of an aTc-inducible promoter, the addition of aTc quenches PhoR activity in medium containing Gly-3P as the sole P source (Fig. 8C). Here, PhoR repression is independent of whether GlpT is present to catalyze the export of intracellular Pi released from Gly-3P, that could potentially repress PhoR from the periplasmic side if the cytoplasmic membrane (Fig. 8C and 8D) (5, 27, 72). Hence, the activity of this two-component system is repressed by Gly-3P catabolism, in the absence of extracytoplasmic Pi. Third, the activation state of PhoR can be influenced by the expression of proteins that alter the cytoplasmic concentration of Pi. Whereas deletion of the gene encoding the housekeeping PitA transporter increases PhoR activity during grown in high Pi medium (Fig. 9B and S7), ectopic PitA expression represses PhoR activity during grown in low Pi medium (Fig. 9B). This later phenotype is also observed during ectopic expression of Pho89, a Pi transporter from the yeast *S. cerevisiae* (Fig. 9B). This indicates that the PhoR repression results from an increase in the levels of cytoplasmic Pi and not by another property of the native PitA transporter. Complementarily, during growth in high Pi medium, PhoR can be activated by eptopic expression of the Pi exporter YjbB (Fig. 9B and 9C).

Together, the results presented in this study establish that PhoB/PhoR activity is controlled by the status of P in the cytoplasm, thereby allowing cells to implement an effective safeguard mechanism against toxicity arising from excessive cytoplasmic Pi resulting from PhoB/PhoR hyperactivation (1-4, 22, 23, 31, 32). That is, control of PhoB/PhoR by a cytoplasmic signal enables cells to maintain balanced growth by meeting their biosynthetic demands without exceeding their requirements for Pi.

## Materials and Methods

### Bacterial strains, plasmids, oligonucleotides and strains construction

The bacterial strains and plasmids used in this study are listed in Table S5 and oligonucleotide sequences are presented in Table S6. All *Salmonella enterica* serovar Typhimurium strains are derived from strain 14028s (102) and were constructed by lambda Red–mediated recombination (103). Deletions and gene fusions generated via this method were subsequently moved into clean genetic backgrounds via phage P22-mediated transduction as described (104). Bacterial strains used in recombination and transduction experiments were grown in Luria–Bertani (LB) medium at 30°C or 37°C (103, 104). When required, the LB medium was supplemented with ampicillin (100 µg/mL), chloramphenicol (20 µg/mL), kanamycin (50 µg/mL), gentamicin (18 µg/mL) or apramycin (80 µg/mL), tetracyclin (20 µg/mL).

### Construction of *tetRA-ugpBAECQ* insertion

Phusion High-Fidelity DNA Polymerase (New England Biolabs) and primers 2075 and 2076 were used to PCR-amplify the *km^R^-tetRA* fragment from plasmid pBbB2K-GFP (105). The PCR product was inserted into the chromosome of *Salmonella enterica* using lambda Red–mediated recombination (103). The location of the insertion in kanamycin resistant clones was verified by PCR using primer pairs 2077/2078 and 2079/2080. These clones were also tested for the ability to growth on Gly-3P as the sole P source, in the presence and absence of aTc.

### Construction of plasmid pUHE-YjbB

Phusion High-Fidelity DNA Polymerase (New England Biolabs) and primers W3392 and W3393 were used to PCR-amplify *yjbB* from *Salmonella enterica* 14028s chromosome. PCR product was digested with BamHI and PstSI, and ligated into pUHE-21 (106) previously digested with the same enzymes. Construct was verified by Sanger DNA sequencing with primers 146 and 156.

### Construction of plasmid pUHE-PitA

Phusion High-Fidelity DNA Polymerase (New England Biolabs) and primers W3394 and W3395 were used to PCR-amplify *pitA* from *Salmonella enterica* 14028s chromosome. PCR product digested with BamHI and PstSI, and ligated into pUHE-21 (106) previously digested with the same enzymes. Construct was verified by Sanger DNA sequencing with primers 146 and 156.

### Construction of plasmid pUHE-Pho89

Phusion High-Fidelity DNA Polymerase (New England Biolabs) and primers W3831 and W3832 were used to PCR-amplify *pho89* from *Saccharomyces cerevisiae* DY1457 (107) genome, which was subsequently cloned into linearized pUHE-21 (106) using NEBuilder HiFi DNA Assembly Cloning Kit (New England BioLabs). Construct was verified by Sanger DNA sequencing with primers 146 and 156.

### Physiological experiments growth conditions

Physiological experiments were carried out at 37°C with shaking at 250 rpm in MOPS medium (108) supplemented with 22 mM glucose, 5 mM MgSO_4_, an amino acids mixture (1.6 mM of alanine, glycine, leucine, glutamate and serine; 1.2 mM glutamine, isoleucine and valine; 0.8 mM arginine, asparagine, aspartate, lysine, phenylalanine, proline, threonine, and methionine; 0.4 mM histidine and tyrosine, and 0.2 mM cysteine and tryptophan), and the indicated concentration of P source.

For experiments on solid media supplemented with different P-sources, 1.5% (w/v) noble agar (Difco) was added into MOPS minimal medium described above supplemented with 0.5 mM of the indicated P-source. Bacteria were incubated for 18-24 h at 37°C. P-shifting experiments were conducted as follows: after overnight (∼16 to 20 h) growth in MOPS medium containing 1 mM K_2_HPO_4_, cells were inoculated (1:100) in a similar fresh MOPS medium and grown until an optical density at 600 nm (OD_600_) of approximately 0.4. Subsequently, cells were washed three times in MOPS medium lacking K_2_HPO_4_, and resuspended in fresh MOPS medium supplemented with the indicated P-source. Cells were subsequently propagated for the corresponding amount of time. During physiological experiments, selection of plasmids was accomplished by the addition of ampicillin at 100 µg/mL (overnight growth) or 30 µg/mL (experimental condition), chloramphenicol at 20 µg/mL (overnight growth) or 10 µg/mL (experimental condition). Heterologous expression of proteins was achieved by supplementing cultures with the indicated concentrations of isopropyl β-D-1-thio-galactopyranoside (IPTG) and 0.5 µg/mL of anhydrotetracycline (aTc). Unless otherwise stated in figure legends, the following organic phosphorus sources were used at 1 mM: 2-aminoethylphosphonic acid, adenosine 5’-monophosphate disodium salt, adenosine 5’-diphosphate disodium salt dihydrate, adenosine 5’-triphosphate disodium salt, guanosine 5’-triphosphate sodium salt, fructose-6-phosphate disodium salt, glucose-6-phosphate disodium salt, phosphoenol pyruvic acid monopotassium salt, phosphonoacetic acid, *sn*-glycerol 3-phosphate bis(cyclohexylammonium) salt.

### RNA isolation and sequencing

Four independent biological replicates of wild-type and *phoB* strains were grown in MOPS medium supplemented with 1 mM K_2_HPO_4_ to an OD_600_ of 0.4-0.45. Bacterial cultures were subsequently split into two equal fractions of approximately 10 OD_600_ units, and were labeled either as “-P” or “+Pi”. -P-treatment cells were washed twice (6,000 X *g*, 5 min) with MOPS medium lacking K_2_HPO_4_ (or any other P source). Conversely, +Pi-treatment cells were washed in MOPS medium supplemented with 1 mM K_2_HPO_4_. Bacterial cultures were then grown for 35 min to induce the Pi-starvation response. Cultures were then treated with RNAprotect (Qiagen) for 5 min at RT to stabilize mRNA, and then collected by centrifugation (6,000 X *g*, 5 min). Total RNA was isolated by hot-phenol extraction, and further purified using the RNeasy Kit (Qiagen) with on-column DNase I treatment. The eluates were treated with Turbo DNase (Invitrogen) for 30 min before a second round of purification using the RNeasy Kit. rRNA was depleted using the NEBNext® rRNA Depletion kit (Bacteria) and purified from the enzymatic reactions using RNAClean XP magnetic beads (Beckman Coulter) following manufacturer’s instructions. RNA quality and ribosomal depletion was assessed using BioAnalyzer (Agilent). RNA-seq was performed at the Genomics Core Facility (Penn State University). Sequencing mRNA libraries were prepared using Illumina Tru-seq Stranded mRNA library kit. Approximately 5 million 75-nt single reads per sample were determined using a NextSeq High Output (Illumina), with >96% of the reads having a Q30 quality.

### RNA-Seq mapping and differential gene expression analyses

Untrimmed reads were aligned to the *Salmonella enterica* serovar Typhimurium 14028s reference genome (NC_016855.1; NC_016856.1) using the SeqMan NGen (DNASTAR v17.0.2.1). Assembly files were uploaded into Geneious Prime v11 software (109). RPKM, FPKM, and TPM read counts were calculated using Geneious prior to differential expression analysis using the DeSeq2 Geneious plugin (110) with a false discovery rate of 0.1. Genes with log 2-fold changes above and adjusted p-values less than 0.01 were considered significantly different. Supplementary Tables 1 and 2 contain gene lists, adjusted p-values, and log2 fold changes for transcriptome comparisons.

### Immunoblot analysis

*cydB-* and *phoN2*-HA-tagged *Salmonella* cells were downshifted to P-lacking MOPS medium for 35 min. Alternatively, for the detection of ApeE-HA, ZitB-HA and CydB-HA, cells were grown in MOPS medium with either 50 µM or 1 mM K_2_HPO4 during 18 h. Equivalent amounts of bacterial cells normalized by OD_600_ values were collected, washed with PBS, suspended in 50 mM Tris-HCl pH 8.0, 0.5% SDS sample buffer and 1x protease inhibitor (Roche). Cells were lysed in a Mini-Beadbeater-96 (BioSpec) and insoluble debris was removed by centrifugation (10 min, 10,000 X*g*, 4°C). Protein extracts were quantified using a Rapid Gold BCA Protein Assay Kit (Pierce), mixed with 4X Laemmli sample buffer, and boiled for 5 min. Equal amounts of protein samples within each group were loaded and resolved using 4–12% NuPAGE gels (Life Technologies), and subsequently electro-transferred onto nitrocellulose membrane (iBlot; Life Technologies) following the manufacturer’s protocol. C-terminal HA-tags were detected using Direct-Blot™ HRP anti-DYKDDDDK Tag Antibody (BioLegend). The blots were developed with the SuperSignal West Femto Chemiluminescent system (Pierce) and visualized with an Amersham Imager 600 (GE Healthcare Life Sciences). Mouse-anti RpoB antibody (Thermo Fisher Scientific) was used as the loading control.

### Monitoring gene expression via fluorescence

Following overnight growth, bacteria were washed and diluted in 1:100 in MOPS medium containing the indicated concentrations of P sources, and aliquot as technical replicates or triplicates into black, clear-bottom, 96-well plates (Corning). Two drops of mineral oil were used to seal the wells and prevent evaporation, and cultures were grown at 37° C with auto-mixing in a SpectraMax i3x plate reader (Molecular Devices). At the desired time points, the green fluorescence (excitation 485 nm/emission 535 nm) and OD_600_ from the wells of the plates were read. Fluorescence measurements were normalized by the OD_600_ of the samples.

### Image Acquisition, Analysis, and Manipulation

Plate images were acquired using an Amersham Imager 600 (GE Healthcare Life Sciences). ImageJ software (111) was used to crop the edges, rotate, and adjust the brightness and contrast of the images. These modifications were simultaneously performed across the entire set images to be shown. Noncontiguous sectors from single plates are indicated by dashed lines.

## Data availability

The RNAseq data set is available as GEO submission GSE227715.

## Supporting Information

## Acknowledgments

We are grateful to Beny Spira for critical reading of the manuscript and insightful commnents, and Craig Praul for technical assistance with RNA-Seq.

## Funding and additional information

M.H.P. is partially supported by Grant AI148774 from the NIH and funds from The Pennsylvania State University College of Medicine.

## Supporting Information

**Table S1.** RNA-Seq summary.

**Table S2.** PhoB activated genes.

**Table S3.**
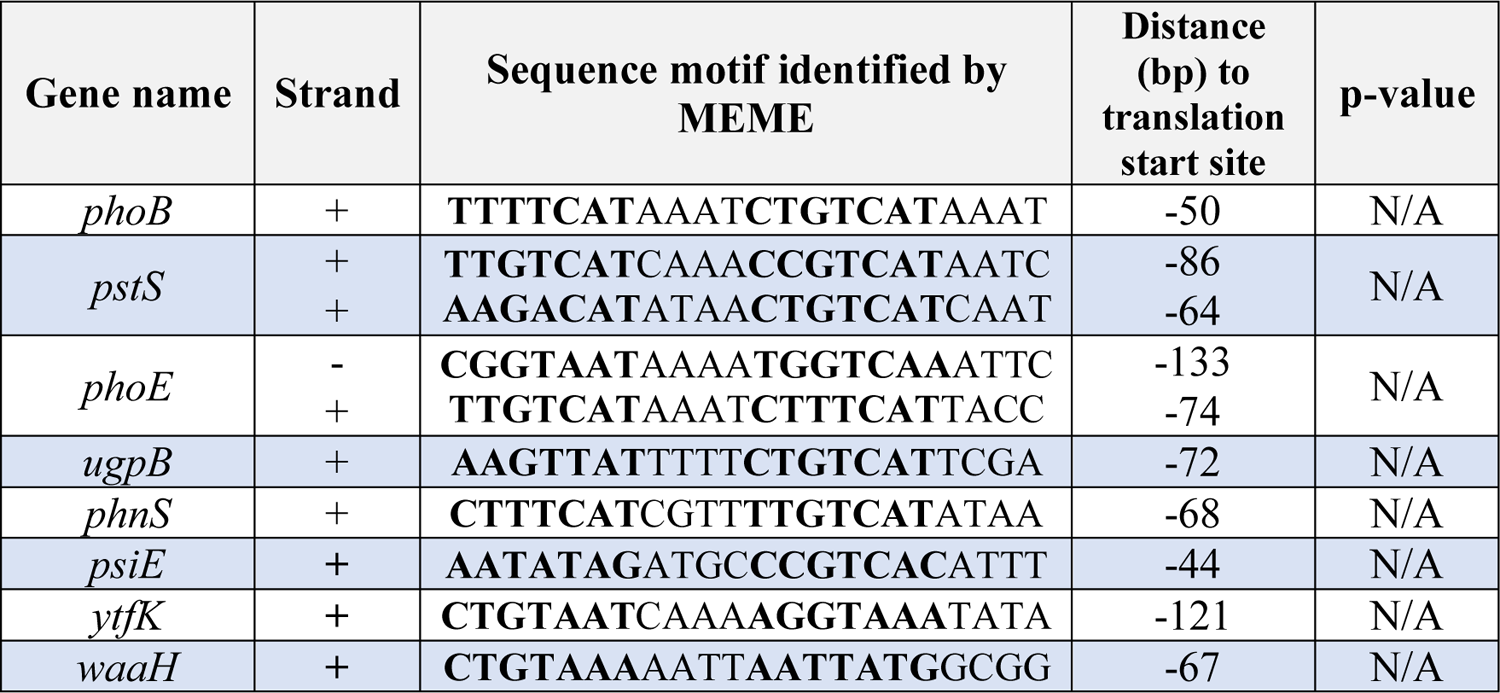
Canonical PhoB-regulated genes used as input for MEME software.

**Table S4.**
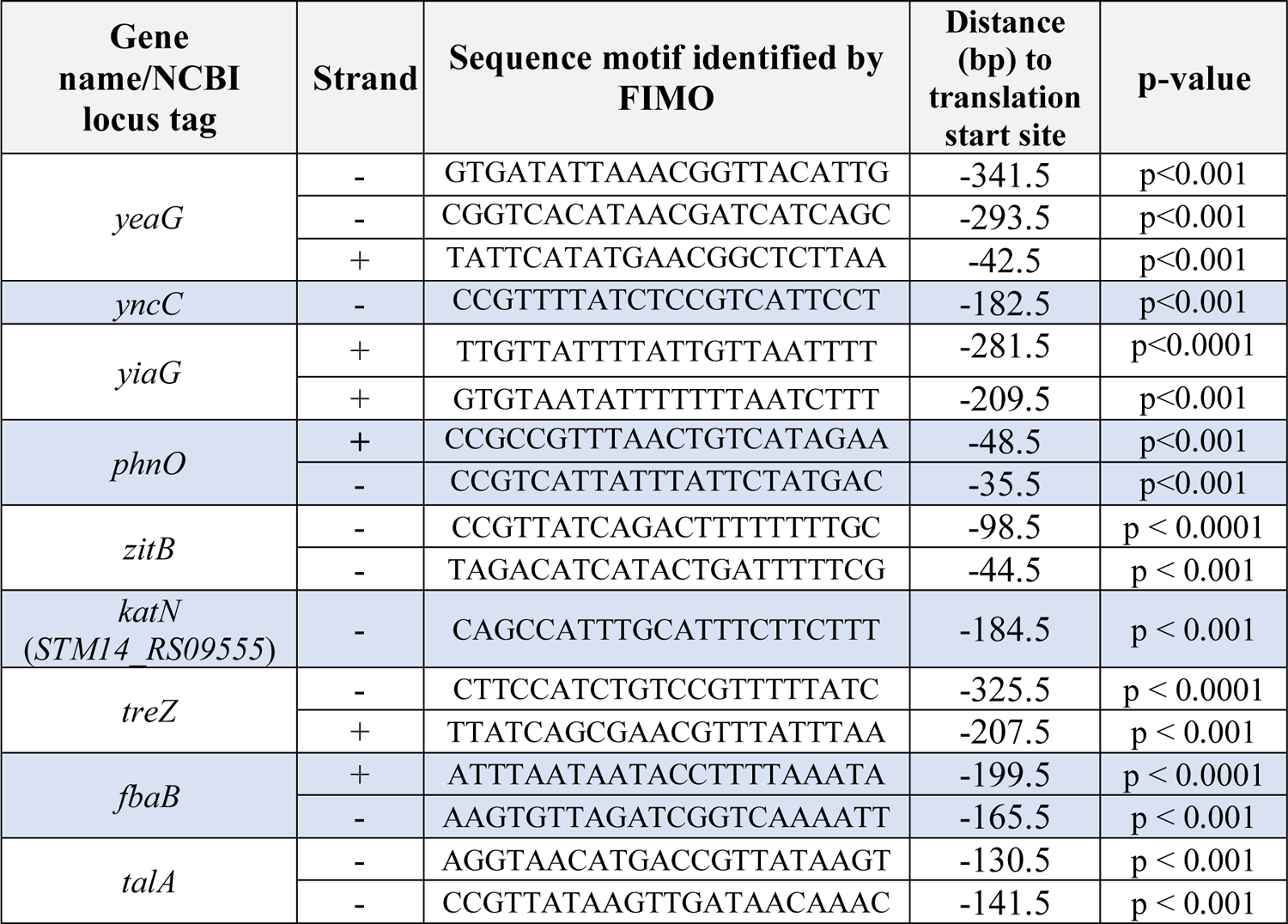
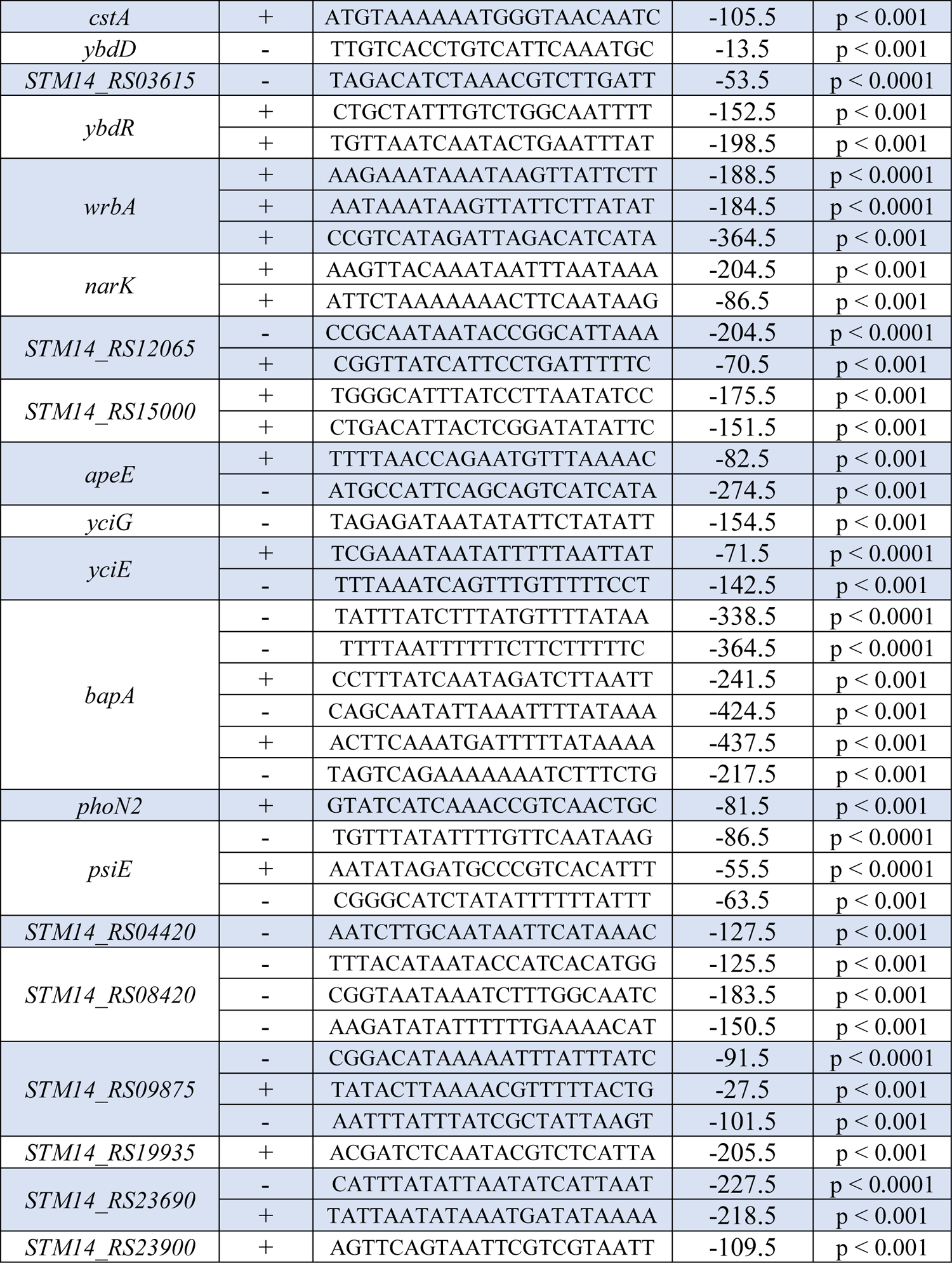
Putative PhoB-motifs predicted by FIMO

**Table S5.**
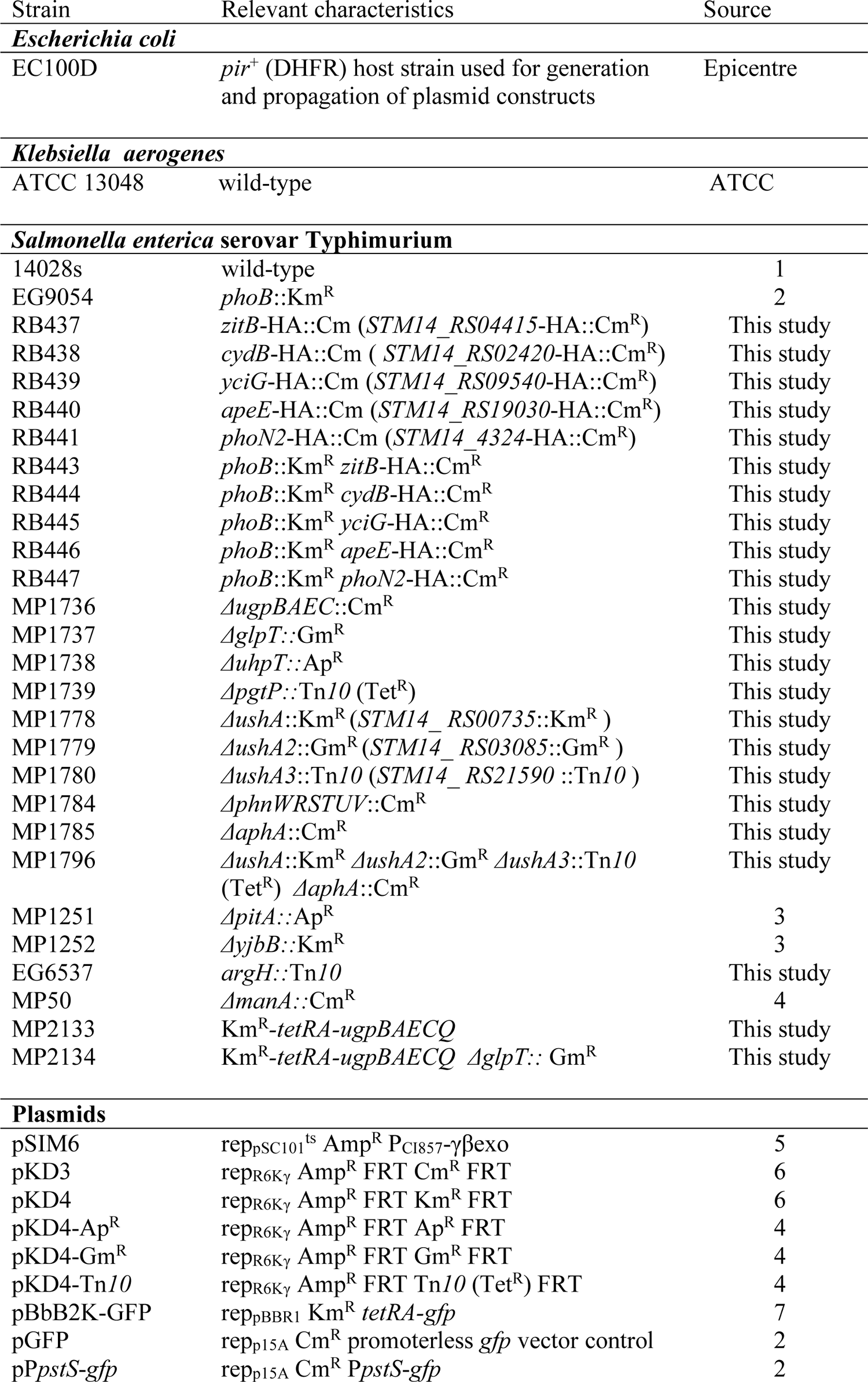
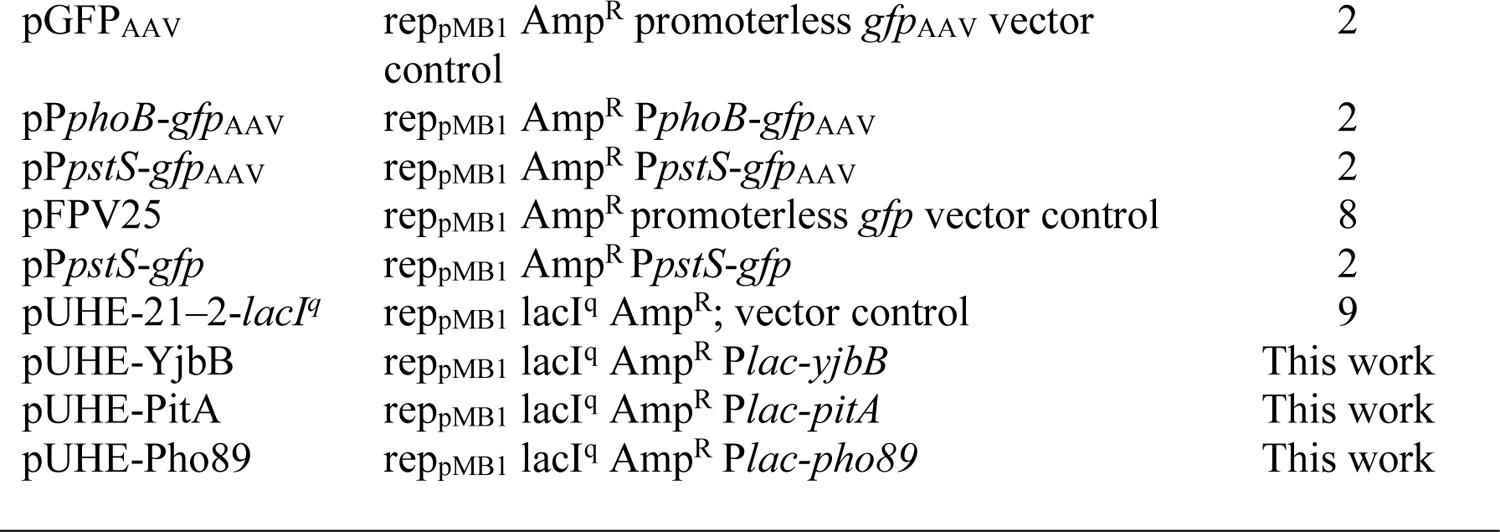
Bacterial strains and plasmids used in this study

**Table S6.**
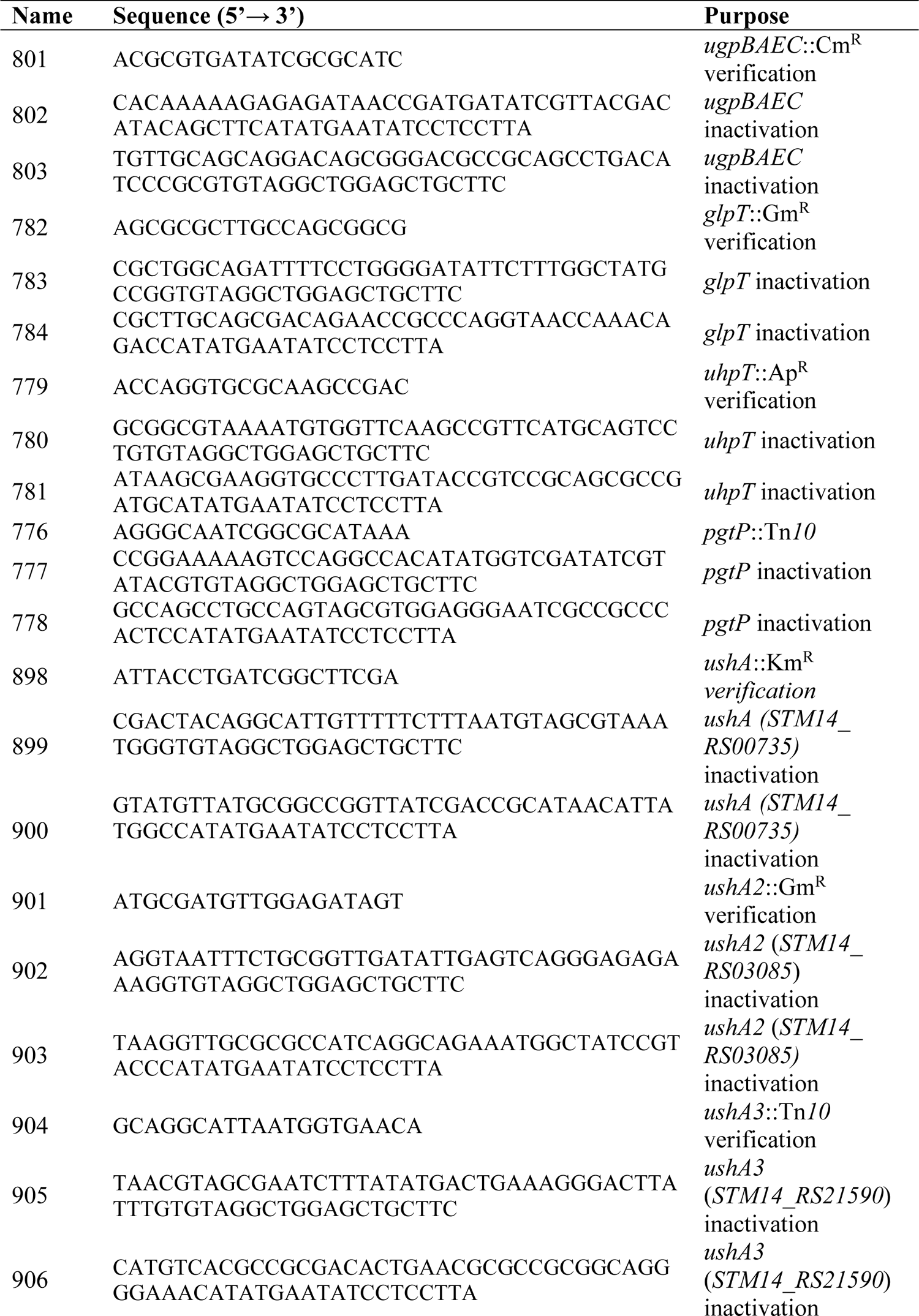
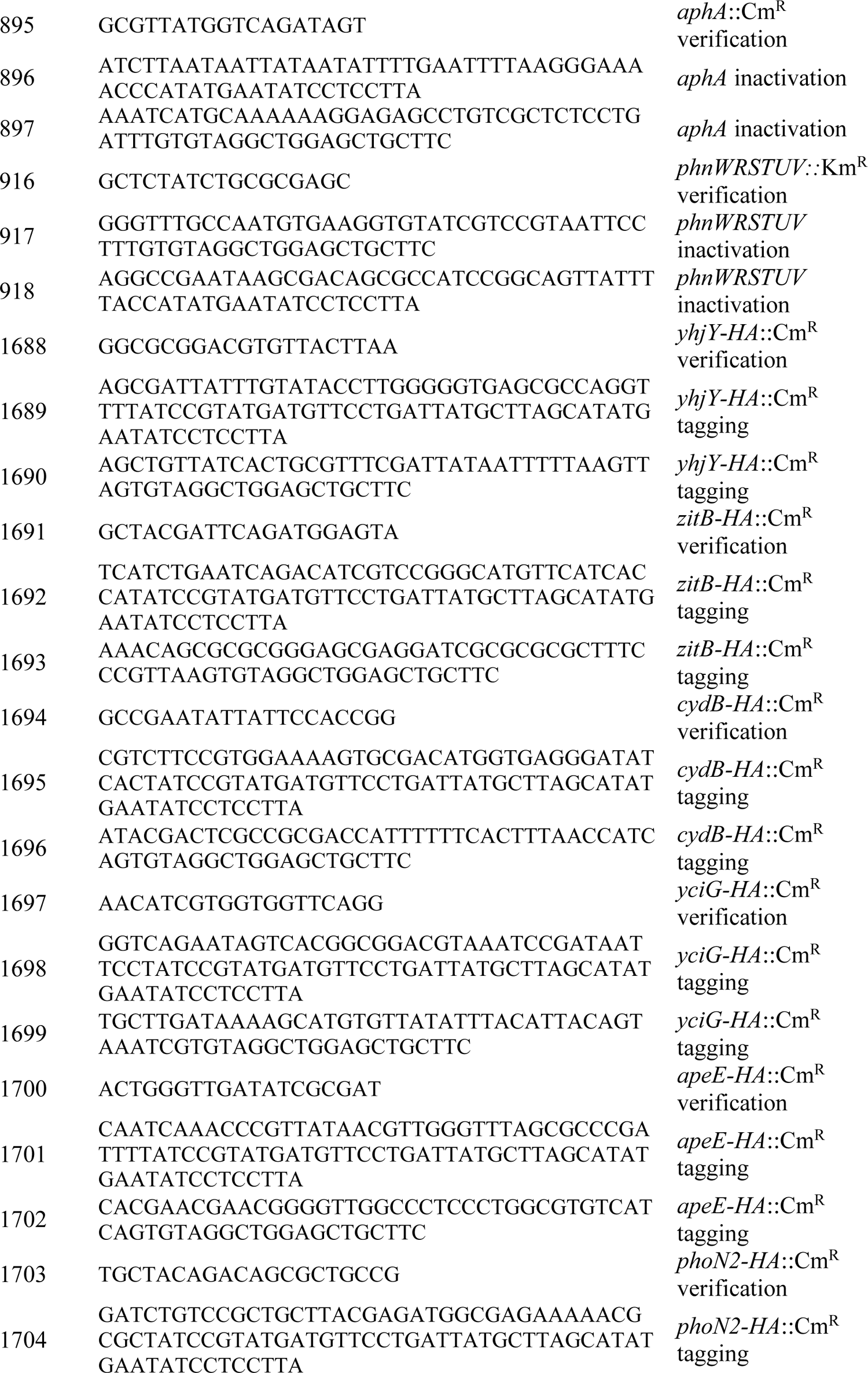
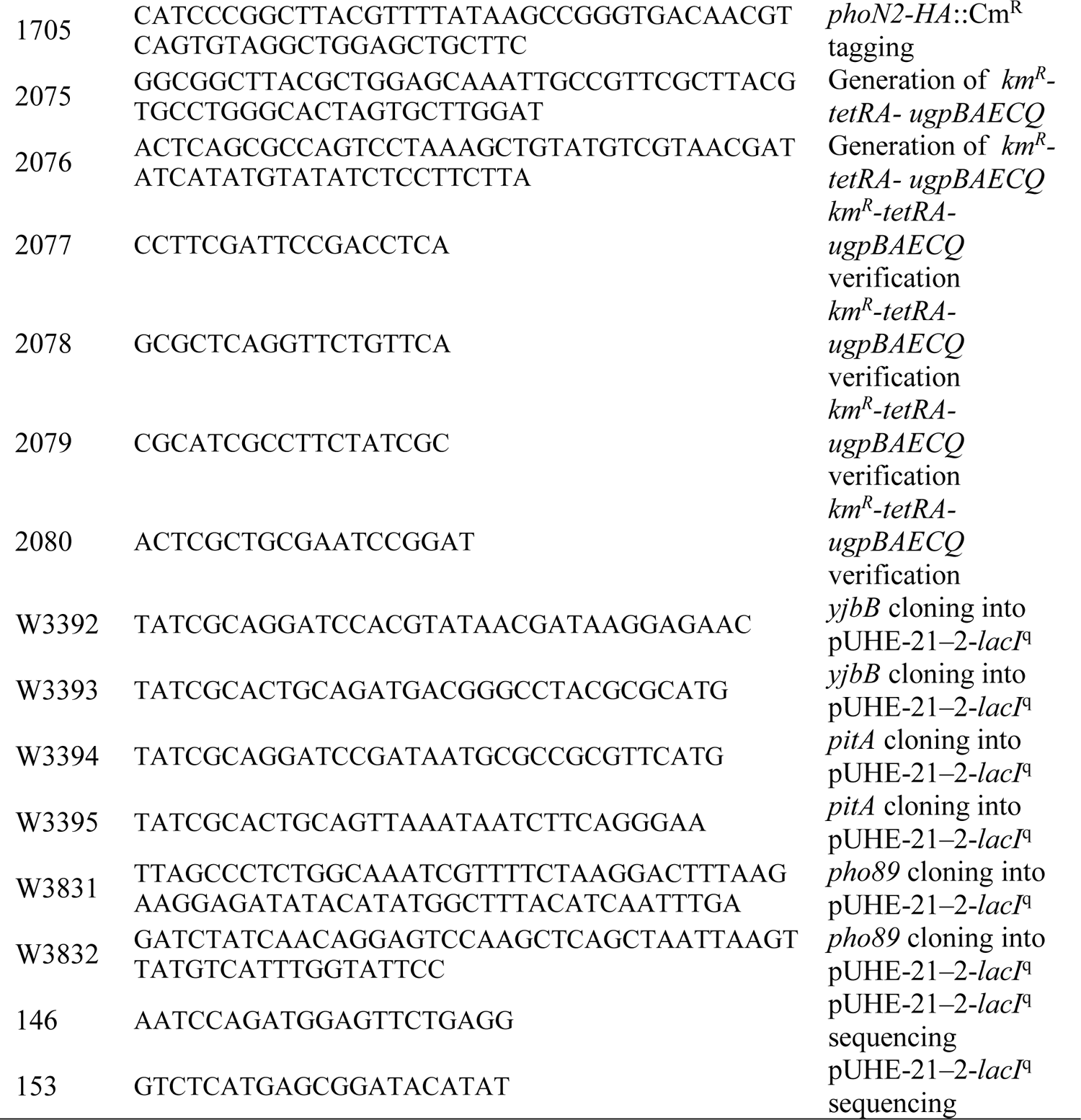
Oligonucleotides sequences used in this study

### Cis regulatory elements of PhoB-regulated genes

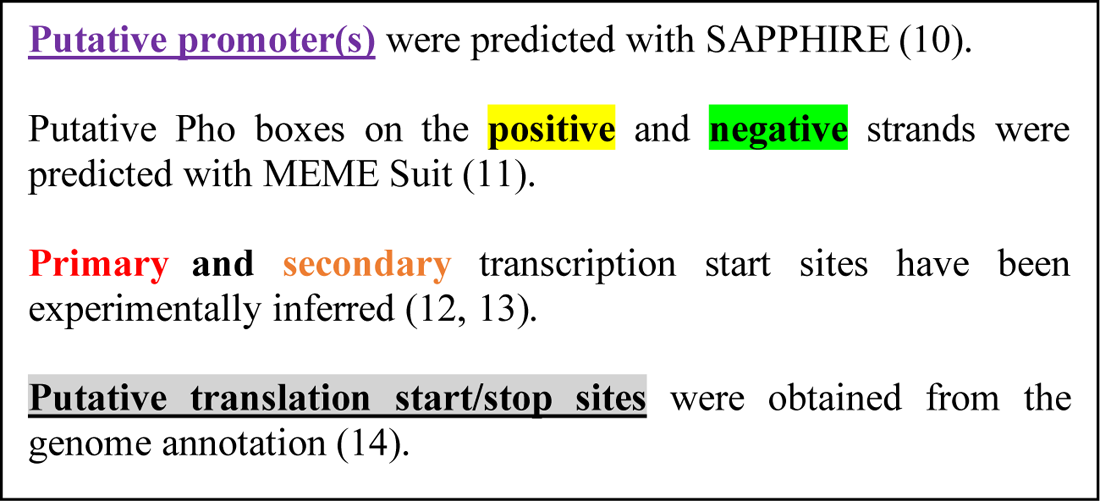

### Canonical PhoB-regulated genes

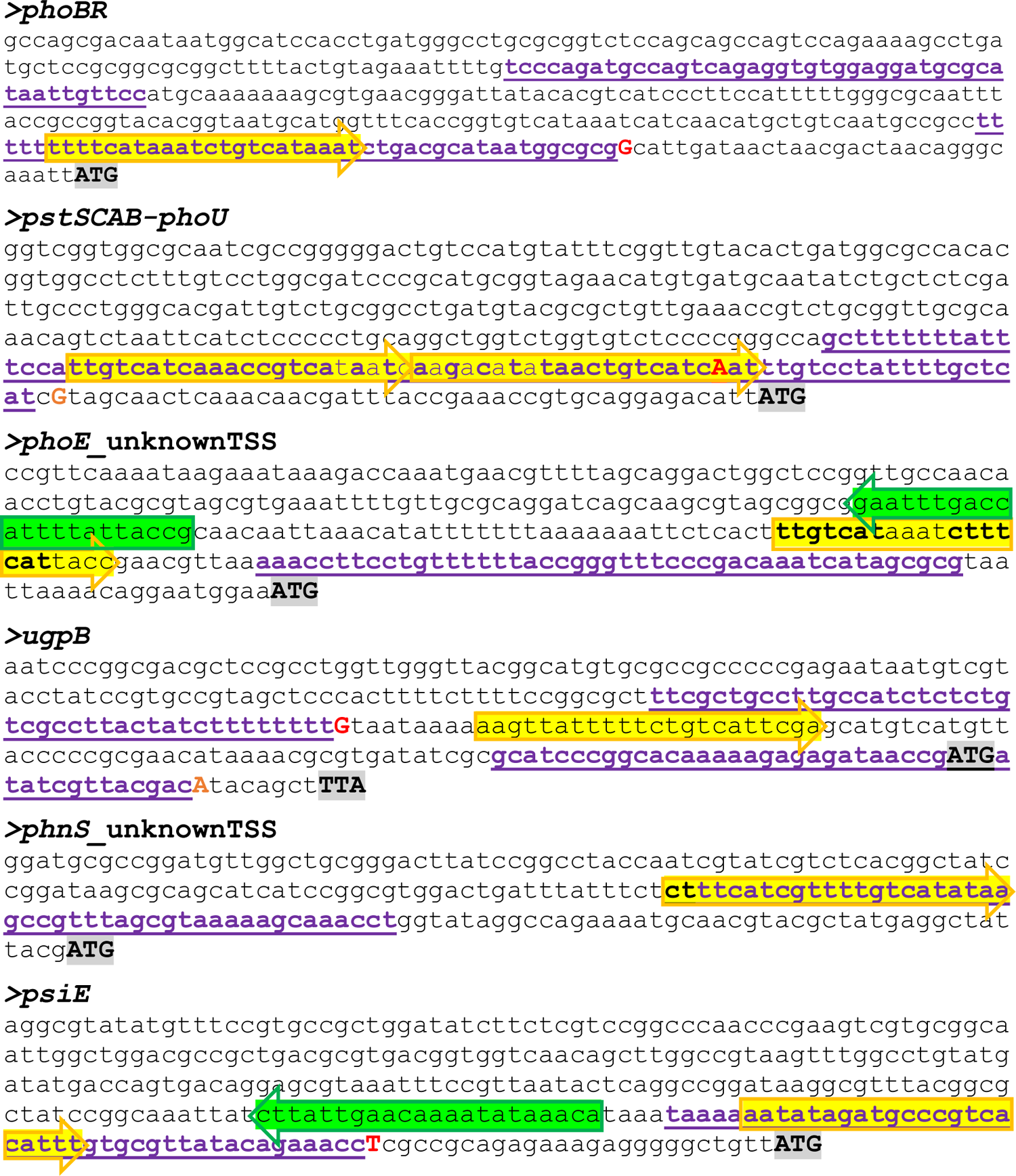
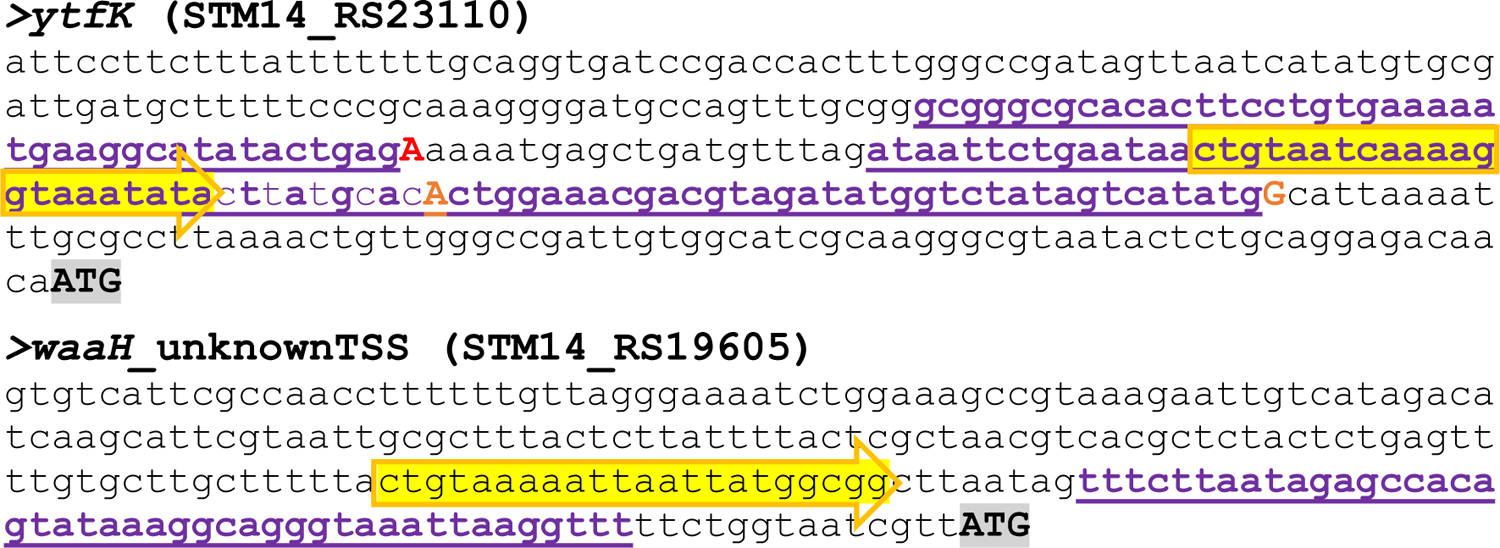

### Putative PhoB-regulated genes

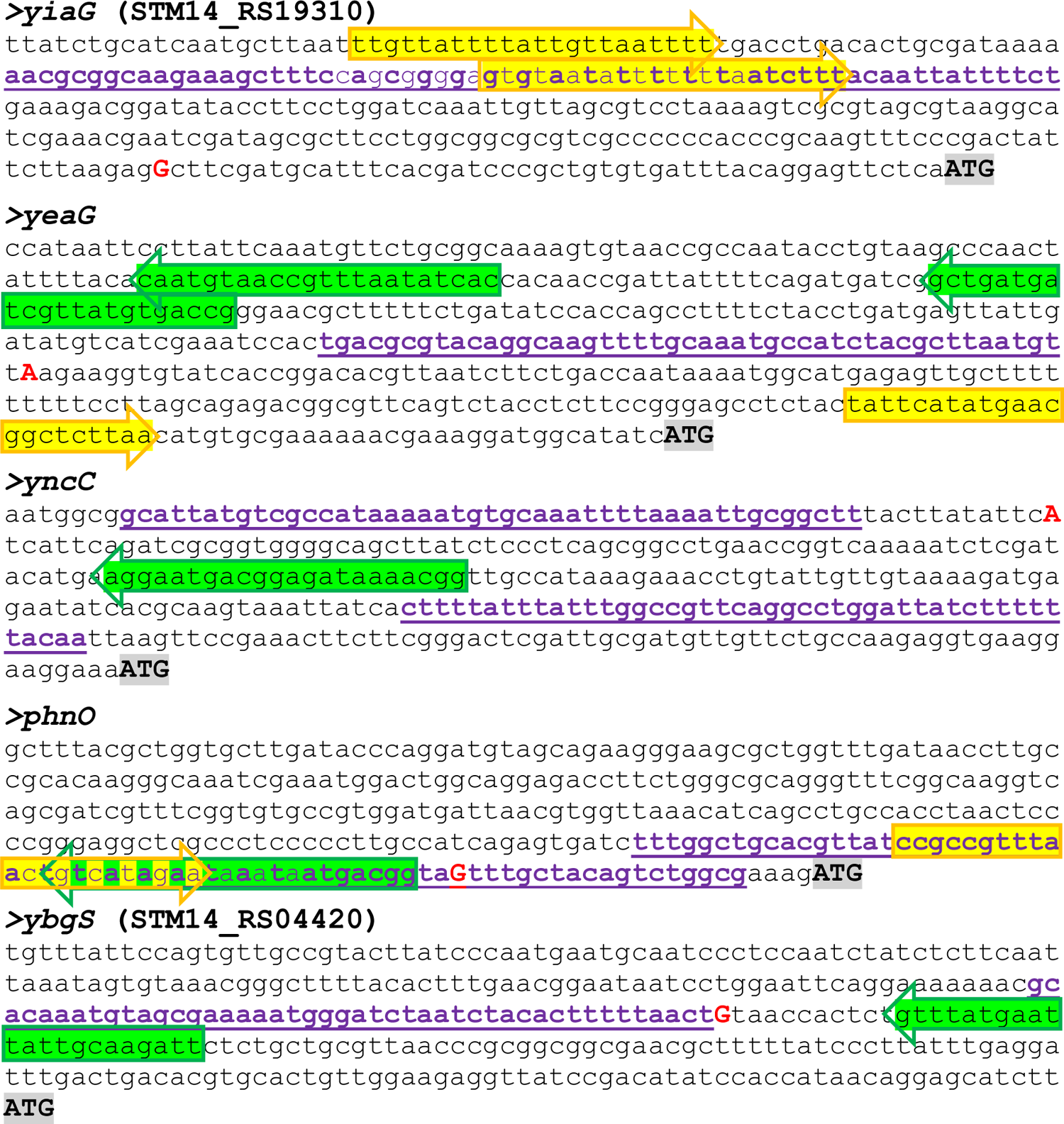
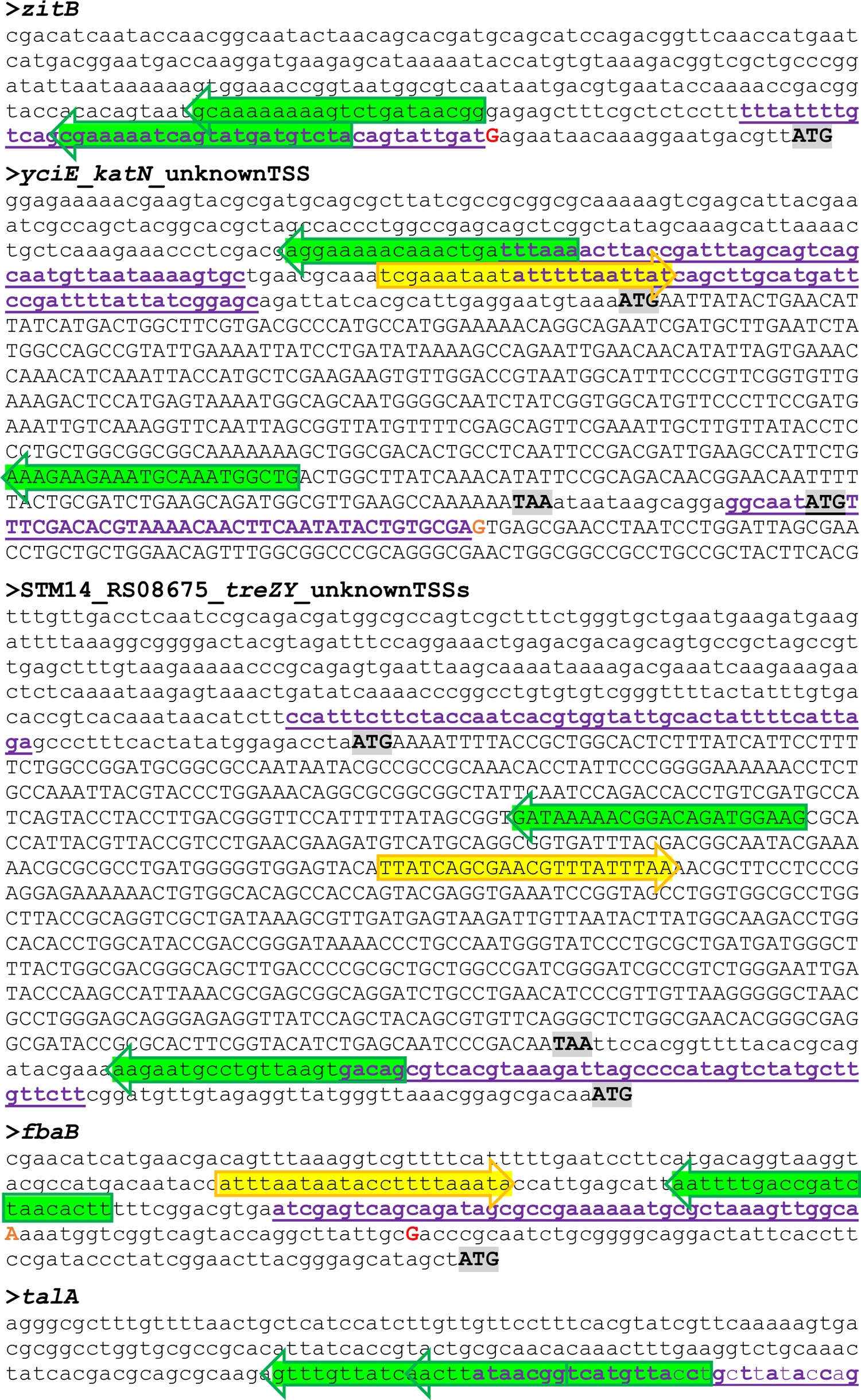
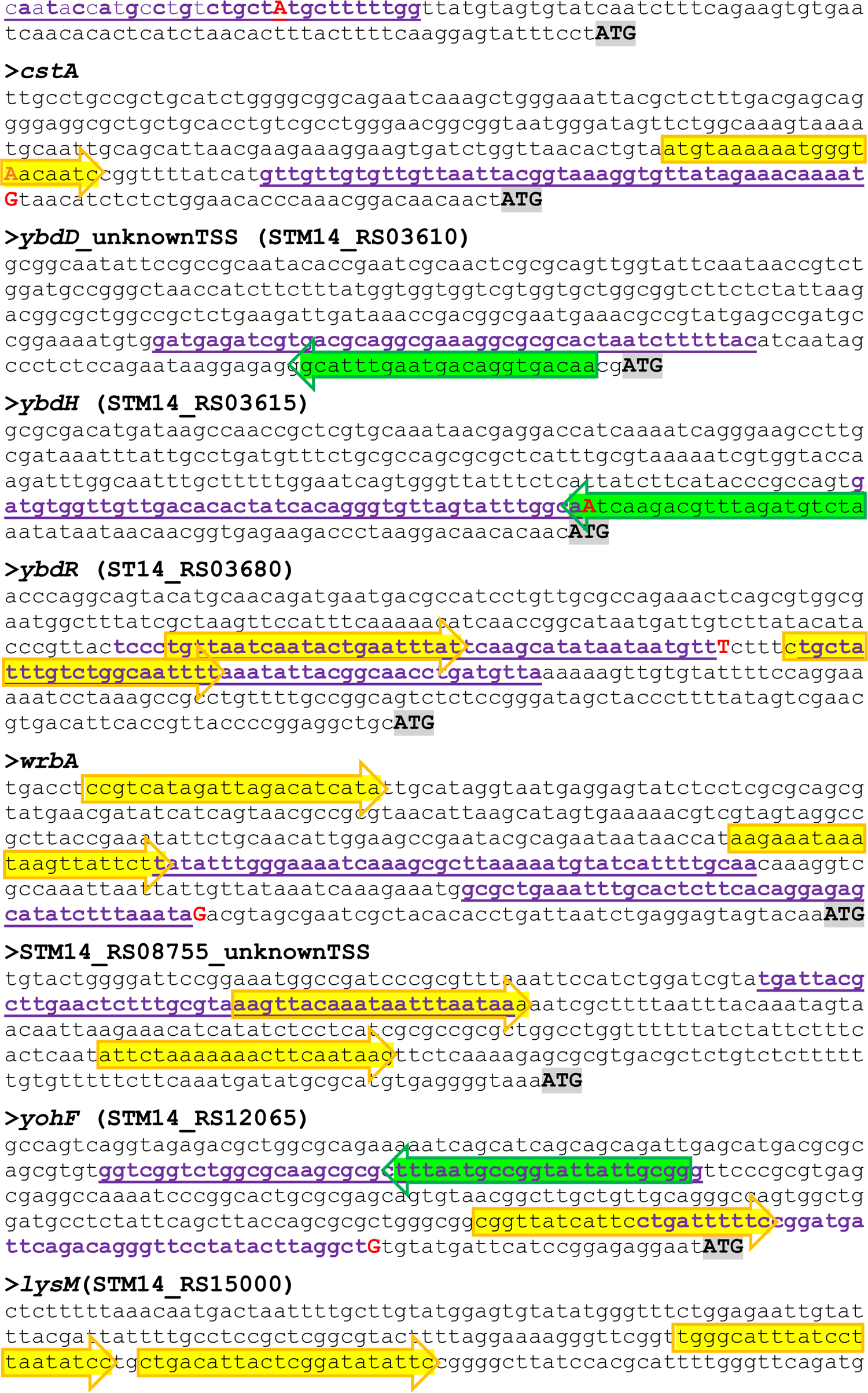
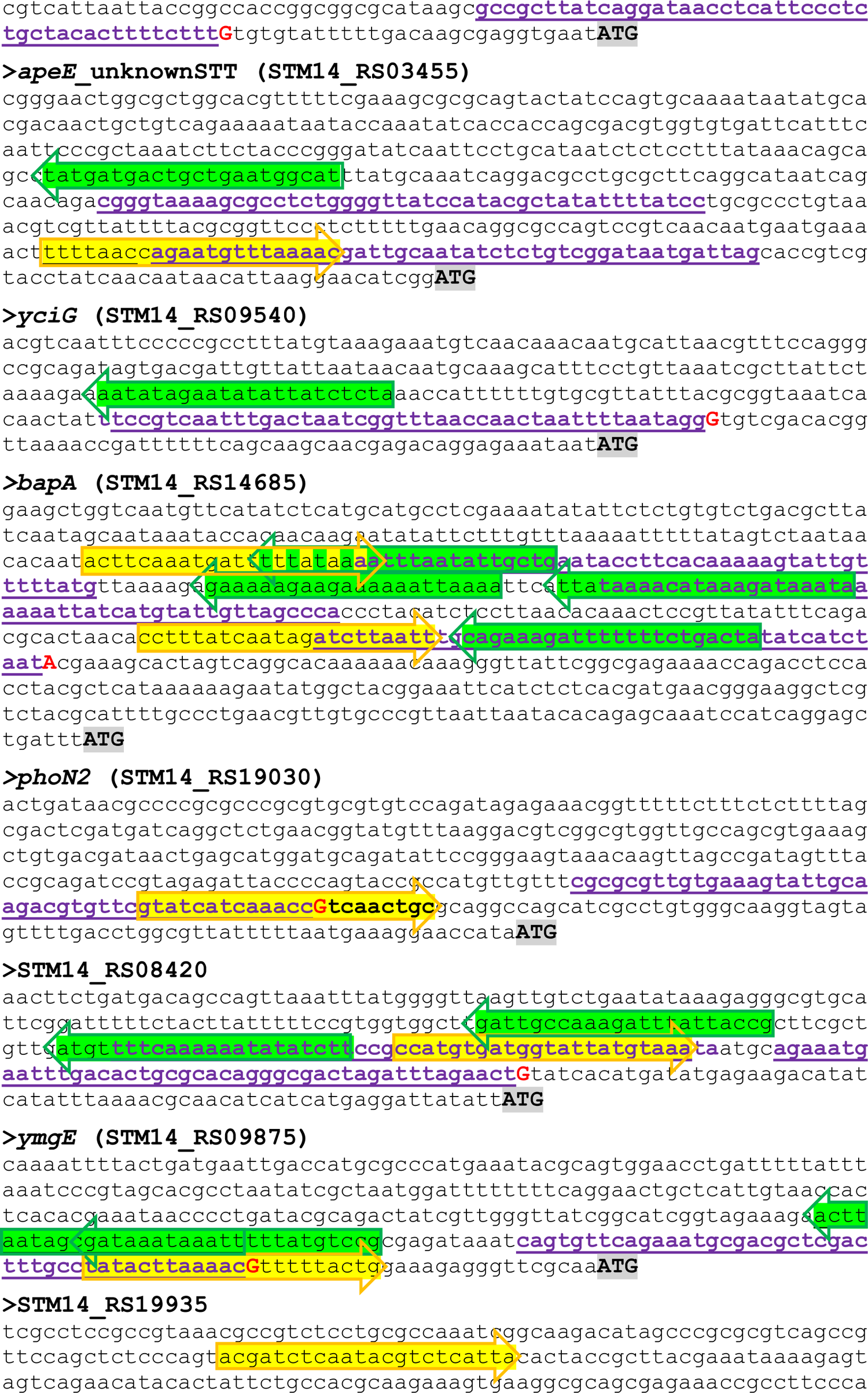
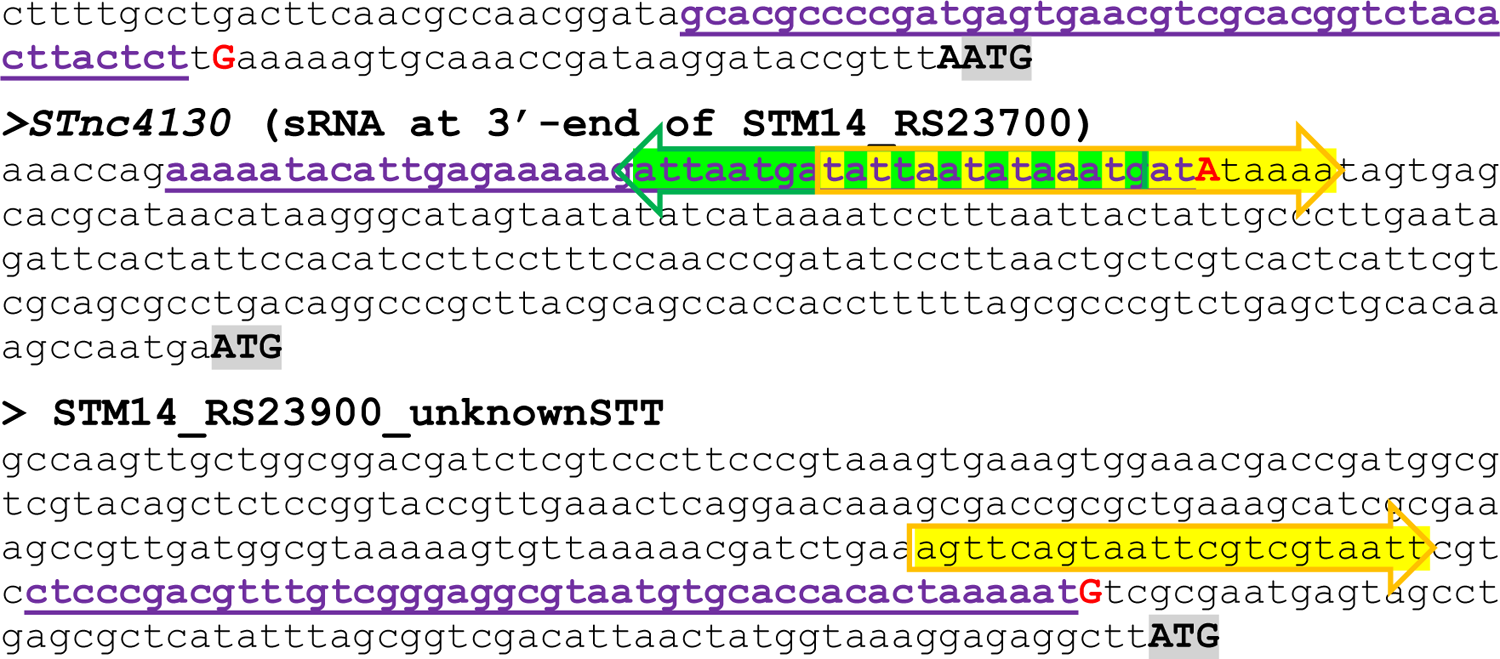

## Supporting Figures

**Figure S1.**
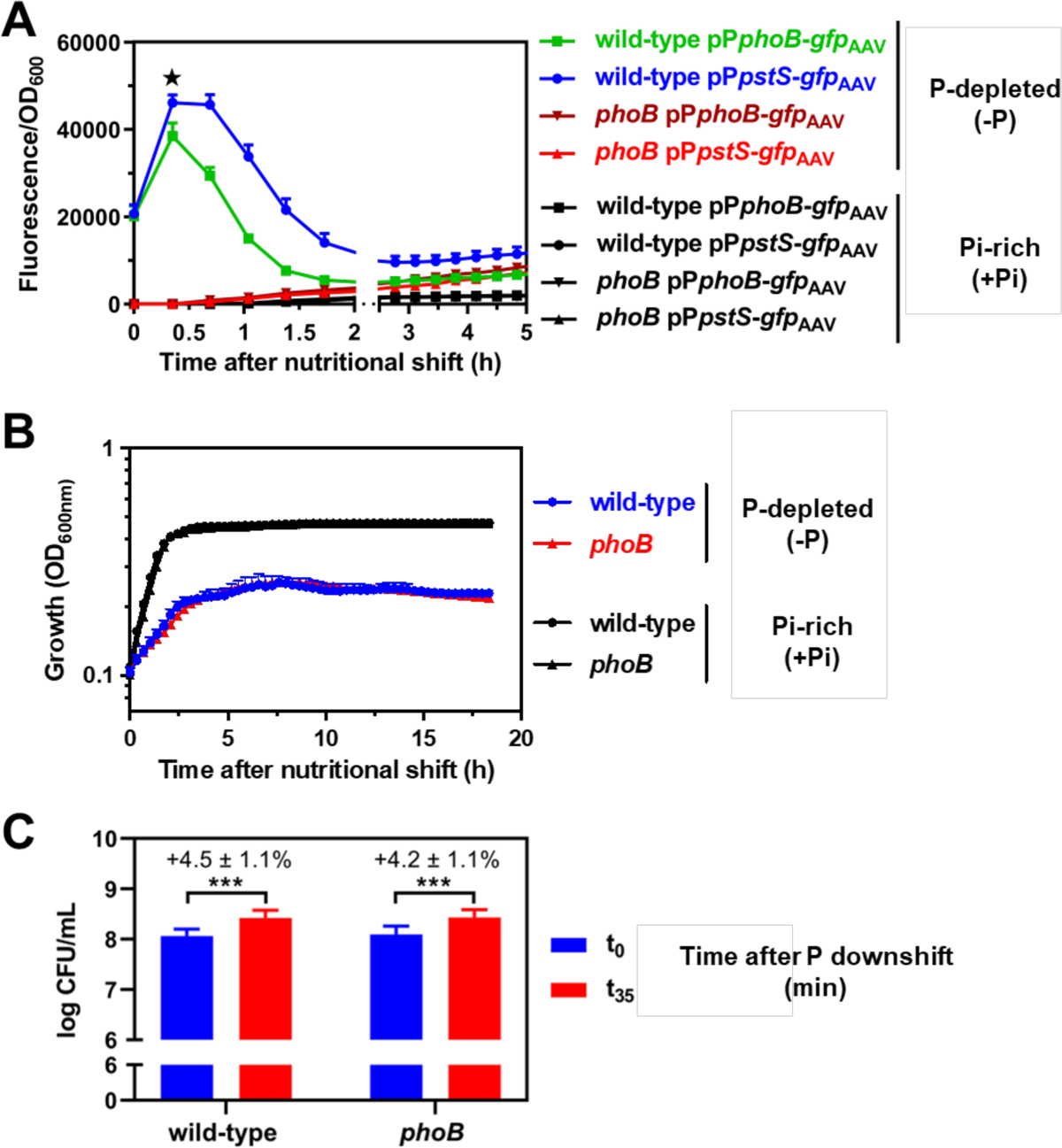
**(A)** Fluorescence from wild-type (14028s) and *phoB* (EG9054) *Salmonella* carrying pP*phoB-gfp*_AAV_ or pP*pstS-gfp*_AAV_. Cultures were propagated to mid-logarithmic phase in MOPS medium containing 1 mM K_2_HPO_4_. Subsequently, cultures were either subjected to a nutritional downshift to MOPS medium lacking K_2_HPO_4_ or any other alternative P source (-P) or maintained in the same medium containing 1 mM K_2_HPO_4_ (+Pi) as control (see Materials and Methods for further experimental details). The maximum GFP expression from *phoB* and *pstS* transcriptional fusions occurs at 35 min following the removal of P, and is indicated with a «. (**B)** Growth curve of wild-type (14028s) or *phoB* (EG9054) *Salmonella* harboring pP*pstS-gfp*_AAV_ subjected to the same nutritional shifts described in (A). **(C)** Viable cell counts of wild-type (14028s) and *phoB* (EG9054) *Salmonella* at the beginning (t_0_) and the end (t_35_) of Pi downshift. *** P < 0.001, paired two-tailed *t* test. In all cases, means ± SDs of at least three independent experiments are shown.

**Figure S2.**
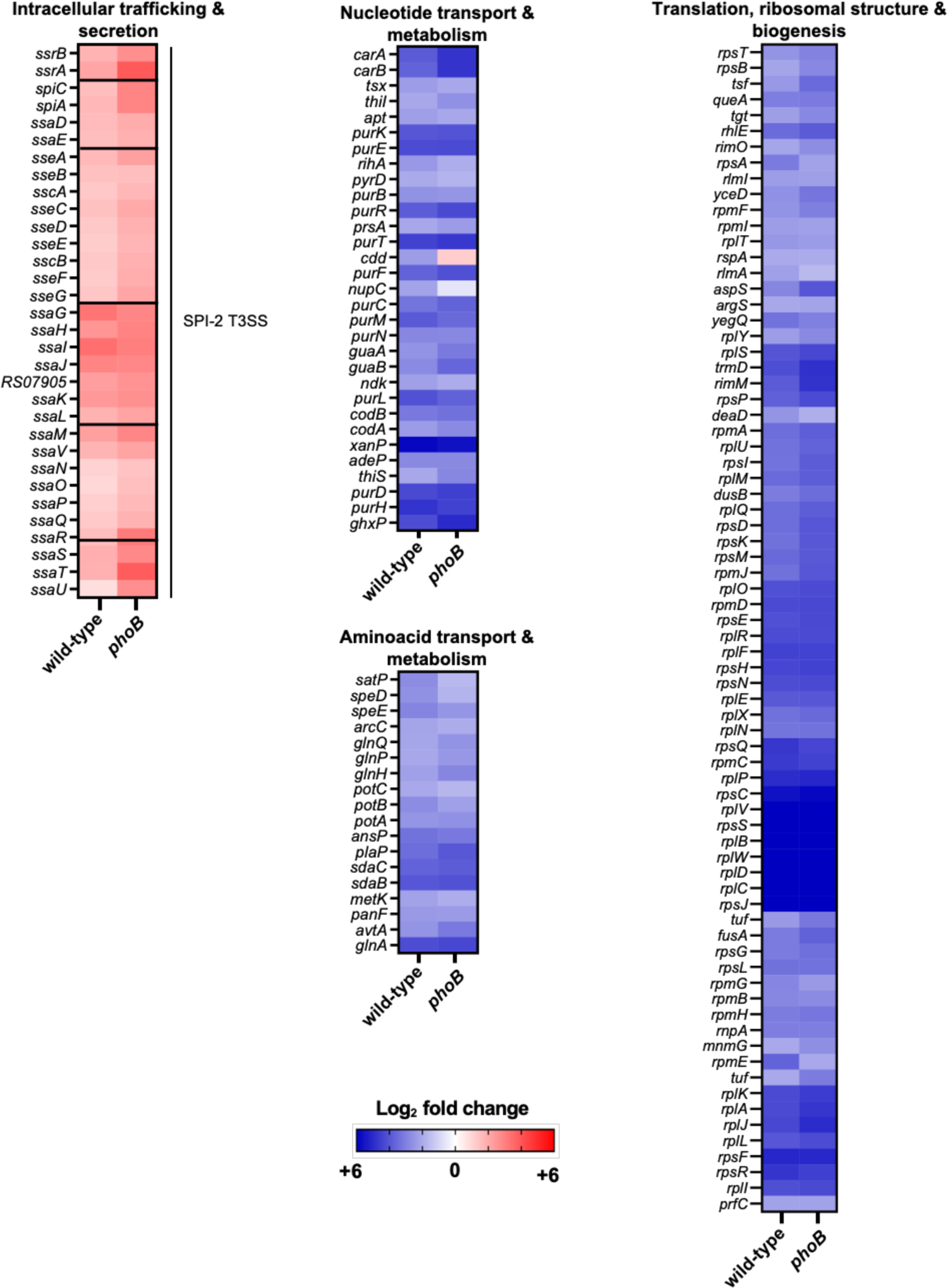
*Salmonella* PhoB-independent response to P starvation. Heatmaps depicting fold changes in transcript levels between -P and +Pi treatments for wild-type (14028s) and *phoB* (EG9054) *Salmonella*. Graphs show selected transcripts from genes that show no significant changes between wild-type and *phoB* during P starvation (Table S1). Displayed genes are organized by COG categories. Note that gene *RS07905* corresponds to NCBI gene locus tag *STM14_RS07905*).

**Figure S3.**
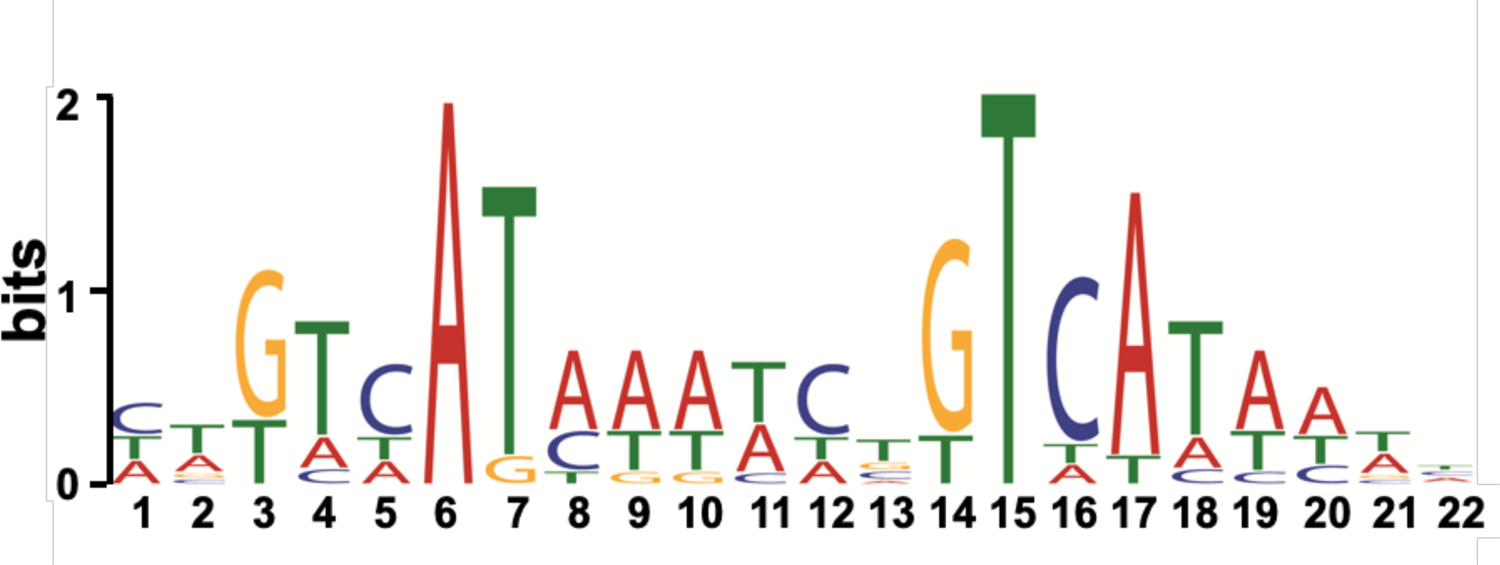
*Salmonella* PhoB binding motif. Sequence logo of the PhoB binding sites. The logo was generated by the MEME algorithm using as input 450-bp upstream of the translation start site of the canonical PhoB-activated operons listed in Table S3 (6).

**Figure S4.**
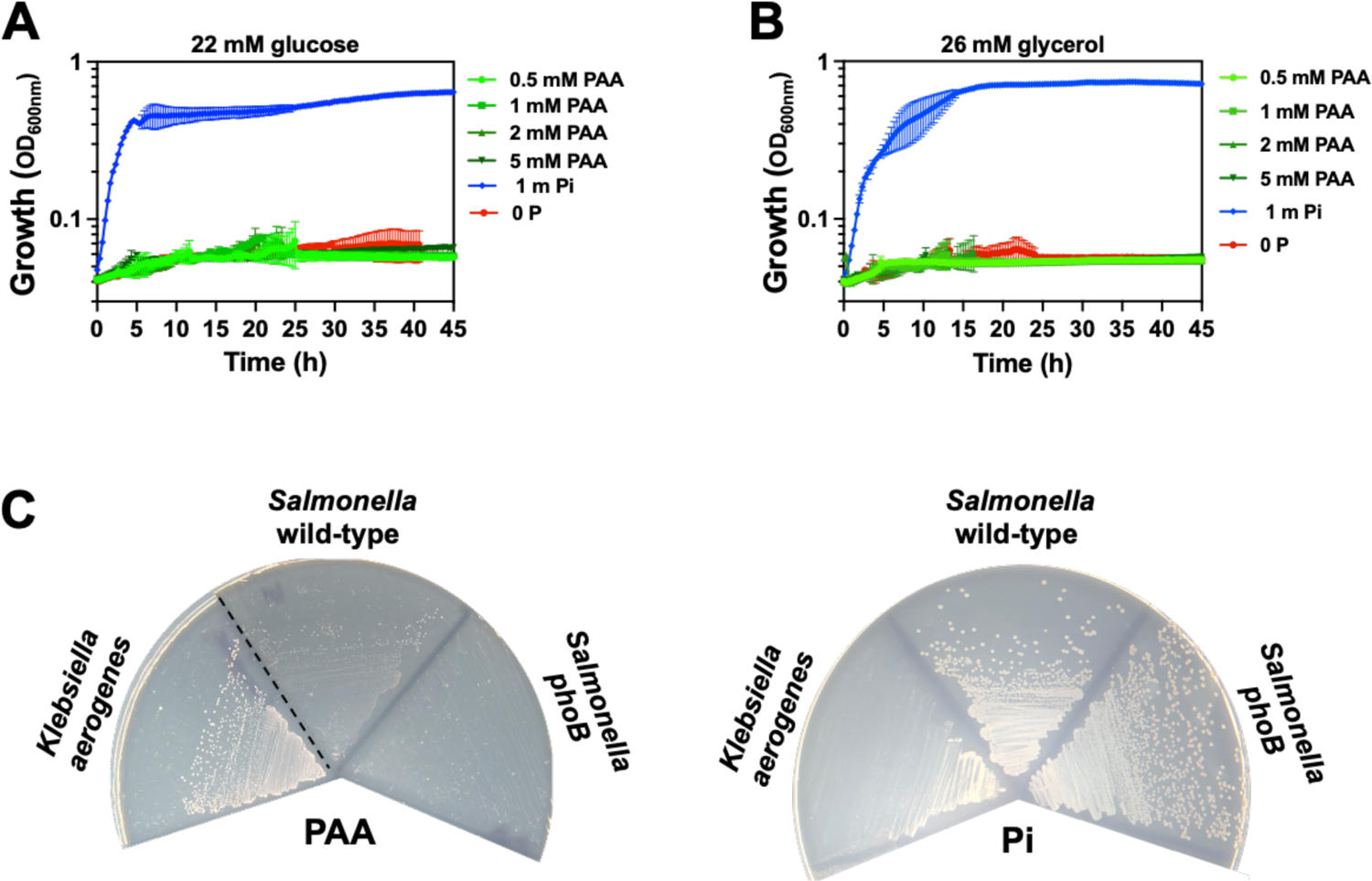
Utilization of phosphonoacetic acid as sole P source. Growth of wild-type *Salmonella* (14028s) in MOPS liquid medium containing (**A**) 22 mM glucose or (**B**) 26 mM glycerol as carbon source and either no P, 1 mM Pi (K_2_HPO_4_) or the indicated concentration of phosphonoacetic acid (PAA) and sole P source. Growth curves show the means ± SDs of three independent biological replicates and are representative of two independent experiments. (**C**) Growth of wild-type *Klebisiella aerogenes* (ATCC13048), and wild-type (14028s) and *phoB* (EG9054) strains of *Salmonella* on MOPS-glucose-noble agar plate containing 1 mM PAA (left) or 1 mM Pi (right) as the sole P source. Plates were incubated at 37°C during 14-18 h before being imaged. Images are representative of three independent experiments. Dashed lines separate non-contiguous sections of the same plate. Note the presence of P-compound(s) in the noble agar that allows the formation of small *Salmonella* colonies. This is not observed during growth in liquid medium (A).

**Figure S5.**
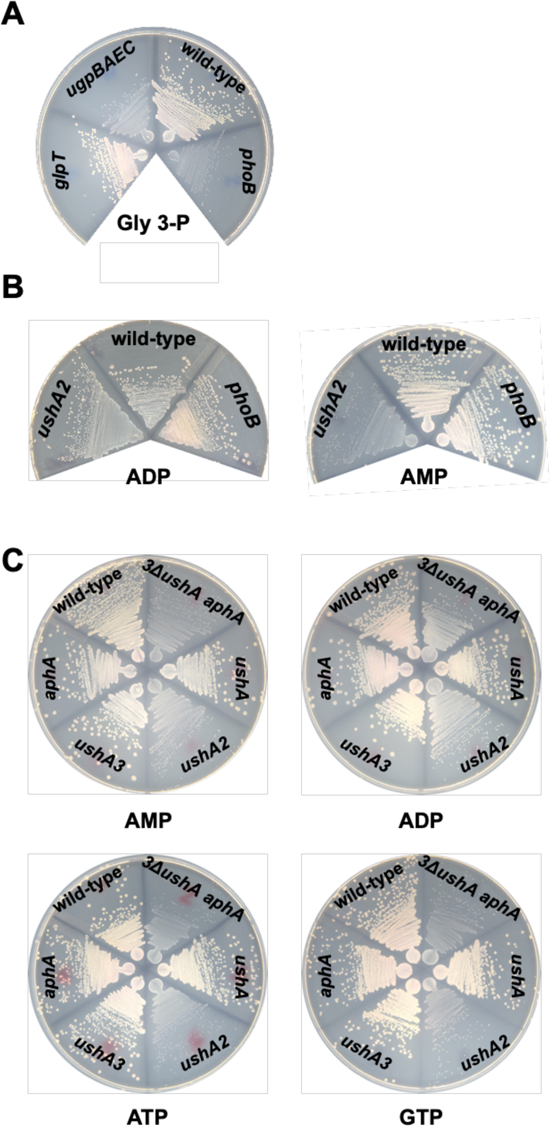
*Salmonella* genetic requirements for the utilization of glycerol-3-phosphate and nucleotides as the only P source. **(A)** Growth of wild-type, *glpT* (MP1735), *ugpBAEC* (MP1736) and *phoB* (EG9054) *Salmonella* strains on MOPS-glucose-noble agar plate containing 0.5 mM of *sn*-glycerol-3-phosphate (Gly 3-P) as the sole P source. **(B)** Growth of wild-type, *ushA2* (MP1779), and *phoB* (EG9054) *Salmonella* on MOPS-glucose-noble agar plates containing 0.5 mM of ADP or AMP as sole P source. **(C)** Growth of wild-type, *ushA* (MP1778), *ushA2* (MP1779), *ushA3* (MP1780), *aphA* (MP1785) and *ushA ushA2 ushA3 aphA* (*3ΔushA aphA*; MP1796) *Salmonella* strains on MOPS-glucose-noble agar plates containing 0.5 mM of either adenosine monophosphate (AMP), adenosine diphosphate (ADP), adenosine triphosphate (ATP) or guanosine triphosphate (GTP) as sole P source. In all experiments, plates were incubated at 37°C during 16-18 h. Images are representative of two independent experiments.

**Figure S6.**
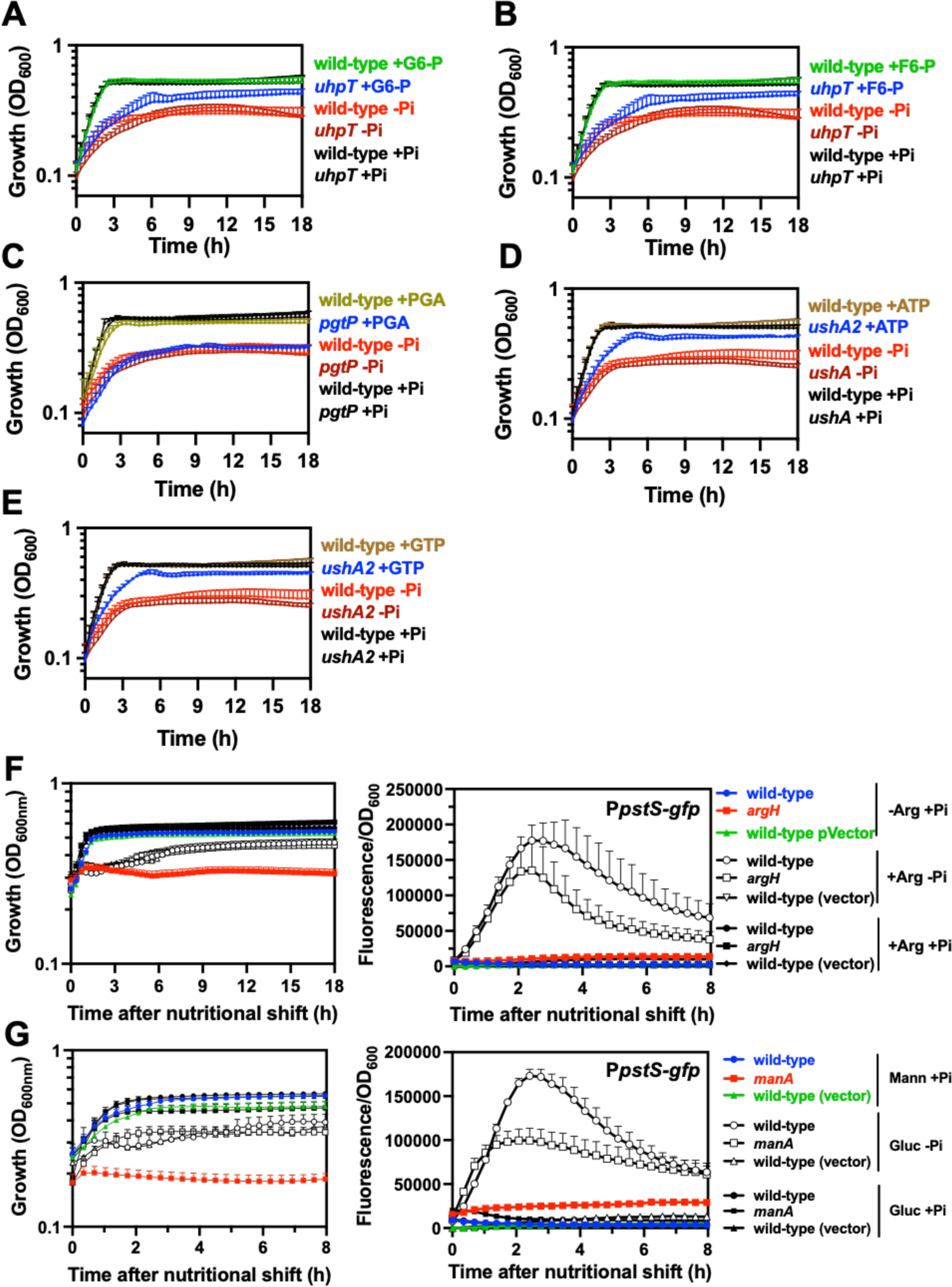
Effect of growth rate on the activity of the P*pstS-gfp* transcriptional fusion. **(A-E)** Growth from wild-type (14028s), *uhpT* (MP1738), *pgtP* (MP1739) and *ushA2* (MP1779) *Salmonella* carrying pP*pstS-gfp*. Growth curves are derived from the experiments outlined in Fig. 6. Prior to performing the readings, cultures were grown to mid-logarithmic phase in MOPS medium containing 1 mM Pi (K_2_HPO_4_), washed in MOPS medium lacking a P source and resuspended in MOPS medium containing 1 mM of the indicated organic P source (G6-P: glucose 6-P; F6-P: fructose 6-P; PGA: 3-phosphoglyceric acid; ATP: adenosine triphosphate), 1 mM Pi (K_2_HPO_4_) or lacking P. Means ± SDs of at least three independent experiments are shown. **(F)** (Left) growth curve and (right) corresponding fluorescence of wild-type (14028s) and *argH* (EG6537) *Salmonella* harboring pP*pstS-gfp* or the vector control (pVector, pFPV25). Cultures were grown to mid-logarithmic phase in MOPS medium containing 1.6 mM arginine and 1 mM Pi (K_2_HPO_4_), washed in MOPS medium lacking arginine and a P source. Cells were subsequently resuspended in either MOPS medium containing or lacking 1.6 mM arginine (+Arg; -Arg) and 1 mM K_2_HPO_4_ (+Pi; -Pi). Growth and green fluorescence were then monitored for 8 h. **(G)** (Left) growth curve and (right) corresponding fluorescence of wild-type (14028s) and *manA* (MP50) *Salmonella* harboring pP*pstS-gfp* or the vector control (pVector, pFPV25). Cultures were grown to mid-logarithmic phase in MOPS medium containing 22 mM glucose and 1 mM K_2_HPO_4_. Cells were washed in MOPS medium lacking carbon and P and then resuspended in MOPS medium containing either 22 mM mannose (+Man) or 22 mM glucose (+Glc) as C sources, and either with or without 1 mM K_2_HPO_4_ (+Pi; -Pi). Growth and green fluorescence were monitored for the next 8 h. Graphs depicted in F-G show the means ± SDs of at least three biological replicates are shown and are representatives of at least 3 experiments.

**Figure S7.**
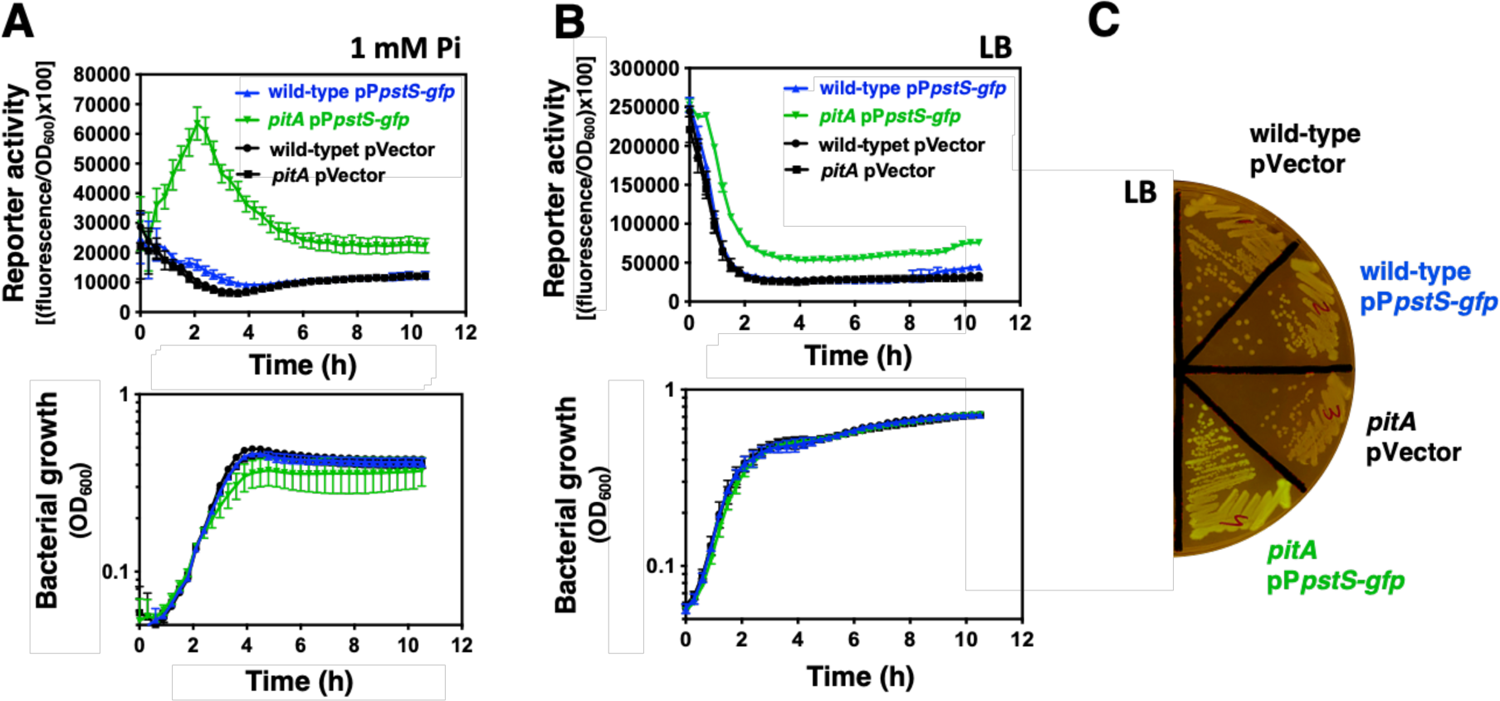
Effect of *pitA* deletion on the activity of the P*pstS-gfp* transcriptional fusion. **(A)** (Top) Fluorescence and (bottom) growth of wild-type (14028s) and *pitA* (MP1251) strains of *Salmonella* harboring pP*pstS-gfp* or the vector control (pVector, pFPV25). Measurements were performed during growth in MOPS glucose medium supplemented with 1 mM Pi (K_2_HPO_4_). **(B)** (Top) Fluorescence and (bottom) growth of strains described in (A). Measurements were performed in LB medium, which contains 2 mM Pi (81). **(C)** Fluorescence of strains described in (A) on LB plates. Plates were incubated at 30°C for 14-16 h. Image is representative of three independent experiments.

